# Genomics of Natural Populations: Gene Conversion Events Reveal Selected Genes within the Inversions of *Drosophila pseudoobscura*

**DOI:** 10.1101/2022.08.15.503618

**Authors:** Stephen W. Schaeffer, Stephen Richards, Zachary L. Fuller

**Affiliations:** Human Genome Sequencing Center, Baylor College of Medicine, One Baylor Plaza, Houston, TX 77030; 23andMe, Inc., Sunnyvale, CA 94086

**Keywords:** Gene Conversion, Inversion, Selection, Local Adaptation, *Drosophila pseudoobscura*

## Abstract

When adaptive phenotypic variation or QTLs map within an inverted segment of a chromosome, researchers often despair because it is thought that the suppression of crossing over will prevent the discovery of selective target genes that contribute to the establishment of the rearrangement. If an inversion polymorphism is old enough, then the accumulation of gene conversion tracts offers the promise that QTLs or selected loci within inversions can be mapped. This study uses the inversion polymorphism of *Drosophila pseudoobscura* as a model system to show that gene conversion analysis is a useful tool for mapping selected loci within inversions. *D. pseudoobscura* has over 30 different chromosomal arrangements on the third chromosome (Muller C) in natural populations and their frequencies vary with changes in environmental habitats. Statistical tests of five *D. pseudoobscura* gene arrangements identified outlier genes within inverted regions based on local clusters of fixed SNP differences. These outlier genes also had potentially heritable variation, either fixed amino acid differences or differential expression patterns among arrangements. Here, we use genome sequences of the inverted third chromosome (Muller C) to infer 98,443 gene conversion tracts for a total coverage of 142 Mb or 7.2 x coverage of the 19.7 Mb chromosome. We estimated gene conversion tract coverage in the 2,668 genes on Muller C and tested whether the number of genes with significantly low coverage was the same for outlier versus non-outlier loci.. Genes with low gene conversion tract coverage were more frequent in the outlier class than the non-outlier class suggesting that selection removes exchanged DNA from the outlier genes more often than non-outlier genes. These data support the hypothesis that the pattern and organization of genetic diversity on the third chromosome in *D. pseudoobscura* is consistent with the capture of locally adapted combinations of alleles prior to inversion mutation events.

## Introduction

Chromosomal inversions were first observed indirectly as recombination modifiers in *Drosophila melanogaster* (Sturtevant 1917) and later demonstrated to be the result of the reordering of genes along chromosomes (Sturtevant 1921). An inversion results from two breaks along a chromosome and the rejoining of the central segment in reverse order. The direct examination of polytene chromosomes from Dipteran larval salivary glands revealed that many species are segregating for inversion polymorphisms on one or more chromosomal arms (Dobzhansky and Sturtevant 1938; Wasserman 1982; Sperlich and Pfriem 1986; Carson 1992; Coluzzi *et al*. 2002). An outstanding question in evolutionary genetics has been whether neutral or selective forces are responsible for establishing inversions in populations (Berdan *et al*. 2023). Quantitative trait loci (QTL) and genes that underlie ecotypes have been found to map within inversions or supergenes in a wide variety of species (Lowry And Willis 2010; Huynh *et al*. 2011; Küpper *et al*. 2015; Mérot *et al*. 2018; Sinclair-Waters *et al*. 2018; Arostegui *et al*. 2019; Ayala *et al*. 2019; Faria *et al*. 2019; Lucek *et al*. 2019; Rennison *et al*. 2019; Han *et al*. 2020; Huang *et al*. 2020; Yan *et al*. 2020; Funk *et al*. 2021; Hager *et al*. 2022; Harringmeyer and Hoekstra 2022). Indirect evidence such as clines, seasonal cycling, and population cage experiments have inferred that inversion polymorphisms have been maintained by selection (Dobzhansky 1944; Wright and Dobzhansky 1946; Dobzhansky 1948; Dobzhansky 1950; Coluzzi *et al*. 1979; Kapun *et al*. 2016; Westram *et al*. 2018). In Drosophila, a number of phenotypic characters have been associated with inversion polymorphisms including mating speed, body size, temperature resistance, wing length, and thorax length (Spiess and Langer 1964; Hoffmann and Weeks 2007; Rego *et al*. 2010). In addition, different karyotypes have differentiation survival under different environmental conditions such as humidity, temperature, food sources (Heuts 1947; Heuts 1948; Cunha 1951; Levine 1952; Battaglia and Smith 1961; Druger 1962; Dolgova *et al*. 2010). What is not clear is what genes lead to the establishment of new chromosomal inversions. Identifying candidate selected genes underlying QTLs and phenotypes within different arrangements have remained elusive because inversions suppress recombination leading to extensive linkage disequilibrium within inverted regions that can mask signatures of selection.

Genomic re-sequencing data have provided insights into the mechanisms for the origin, establishment, and maintenance of chromosomal inversions in natural populations (For a recent review see, Berdan *et al*. 2023). Comparison of sequences between different gene arrangements have allowed the identification of regions of chromosomes where breakage and rejoining events led to the origin of new gene orders (Richards *et al*. 2005; Ranz *et al*. 2007; Harringmeyer and Hoekstra 2022). Ectopic exchange between repeat sequences at breakpoints provide one model for the facilitation of inversion mutations (Fuller *et al*. 2017; Harringmeyer and Hoekstra 2022), while staggered cuts that duplicate genes at breakpoints is an alternative mechanism observed to generate new gene arrangements (Ranz *et al*. 2007; Puerma *et al*. 2016).

Once an inversion mutation arises in the population, it is usually lost because its initial frequency (1/2N) is quite low. On the other hand, a new inversion can be established in the population either due to neutral or selective forces. Like point mutations, a selectively neutral inversion can drift to higher frequency through random genetic drift and over time inter-arrangement divergence will be greatest at the breakpoints because the homogenizing effect of genetic flux among arrangements is greatest in the central regions of the inverted regions (Navarro *et al*. 2000; Guerrero *et al*. 2012). There are several possible mechanisms that can explain how an inversion is established by selection. First, direct or position effects (Sperlich and Pfriem 1986) may create beneficial variation when the breakpoints either disrupt genes, create new hybrid genes, or alter gene expression of boundary genes at or near the lesions. Empirical studies of inversion breakpoints have shown that gene duplications can occur at breakpoints (Ranz *et al*. 2007), but have failed to find evidence for gene disruptions within genes (Fuller *et al*. 2017). Inversion breakpoints can altered gene expression of boundary genes (Puig *et al*. 2002). Recent analyses have shown that chromatin states and architecture may influence where breakpoints occur (Mcbroome *et al*. 2020; Wright and Schaeffer 2022).

A second mechanism proposes that indirect effects associated with the suppression of recombination can maintain advantageous haplotypes within the inverted region leading to elevated linkage disequilibrium among multiple genes within the inverted segment (Kirkpatrick and Barton 2006). These models assume that the new inversion captures sets of beneficial alleles (Charlesworth and Charlesworth 1973). Sets of alleles may be beneficial if they are free of deleterious recessive alleles (Nei *et al*. 1967) or are alleles that contribute to local adaptation (Kirkpatrick and Barton 2006). In the case of the local adaptation model, inversions arise to counter the formation of maladaptive genotypes formed through recombination of alternatively selected chromosomes from different habitats. This typically occurs when the migration parameter (*Nm*) is > 1 and populations exist in a heterogeneous environment (Dobzhansky 1944; Kennington *et al*. 2006; Schaeffer 2008; Cheng *et al*. 2012; Kennington and Hoffmann 2013; Kapun *et al*. 2016). The size of an inversion may also influence whether it is established based on the number of genes involved in local adaptation (Kirkpatrick and Barton 2006; Fuller *et al*. 2017). The size of an inversion and its selective role will be a trade off between the number of selected genes versus the ability for double cross overs to reduce associations among selected genes (Caceres *et al*. 1999). Finally, a new gene arrangement may rise to intermediate frequency because one beneficial allele is trapped within the inverted region (Maynard Smith and Haigh 1974) and arrangements can later accumulate additional selected genes.

Once established, the new inversion will act as an isolated subpopulation and accumulate genetic differences from its ancestral arrangement (Navarro *et al*. 2000). Recombination or genetic flux among different gene arrangements will oppose genetic differentiation keeping sequences homogeneous among inversion types (Navarro *et al*. 1997). Genetic flux is initiated by double strand breaks between inverted chromosomes and are either resolved as genetic cross overs or as gene conversion events. Single cross overs within an inverted region of a Drosophila heterokaryotype will generate unbalanced inviable gametes that are selected against leaving only parental gametes (Sturtevant and Beadle 1936). Cross overs are possible if two cross overs occur within the inverted region, but the probability of double cross over events is small unless the size of the inverted segment is large (Navarro *et al*. 1997). Gene conversion events, on the other hand, are genetic exchanges at double strand breaks that are resolved as non-cross over events leading to the exchange of small segments of 200 to 300 nucleotides among arrangements without the formation of unbalanced gametes (Chovnick 1973; Rozas and Aguadé 1993; Rozas and Aguadé 1994; Schaeffer and Anderson 2005) (Figure 1). Conversion happens when the DNA strand from one homologous chromosome (pink strand in Figure 1C) invades the other homolog creating heteroduplexes everywhere where there are SNP differences. Repair of the heteroduplexes in favor of the invading DNA strand leads to the gene conversion event (Figure 1 D). Gene conversion events occur uniformly across the inverted segment, but the homogenizing effect is limited because of the small DNA segment exchanged among chromosomes (Navarro *et al*. 1997; Miller *et al*. 2012; Miller *et al*. 2016; Korunes and Noor 2019). Overall, crossing over is expected to have a greater homogenizing effect than gene conversion because larger genomic segments are exchanged, but crossing over is less likely to occur relative to gene conversion events.

**Figure 1.**
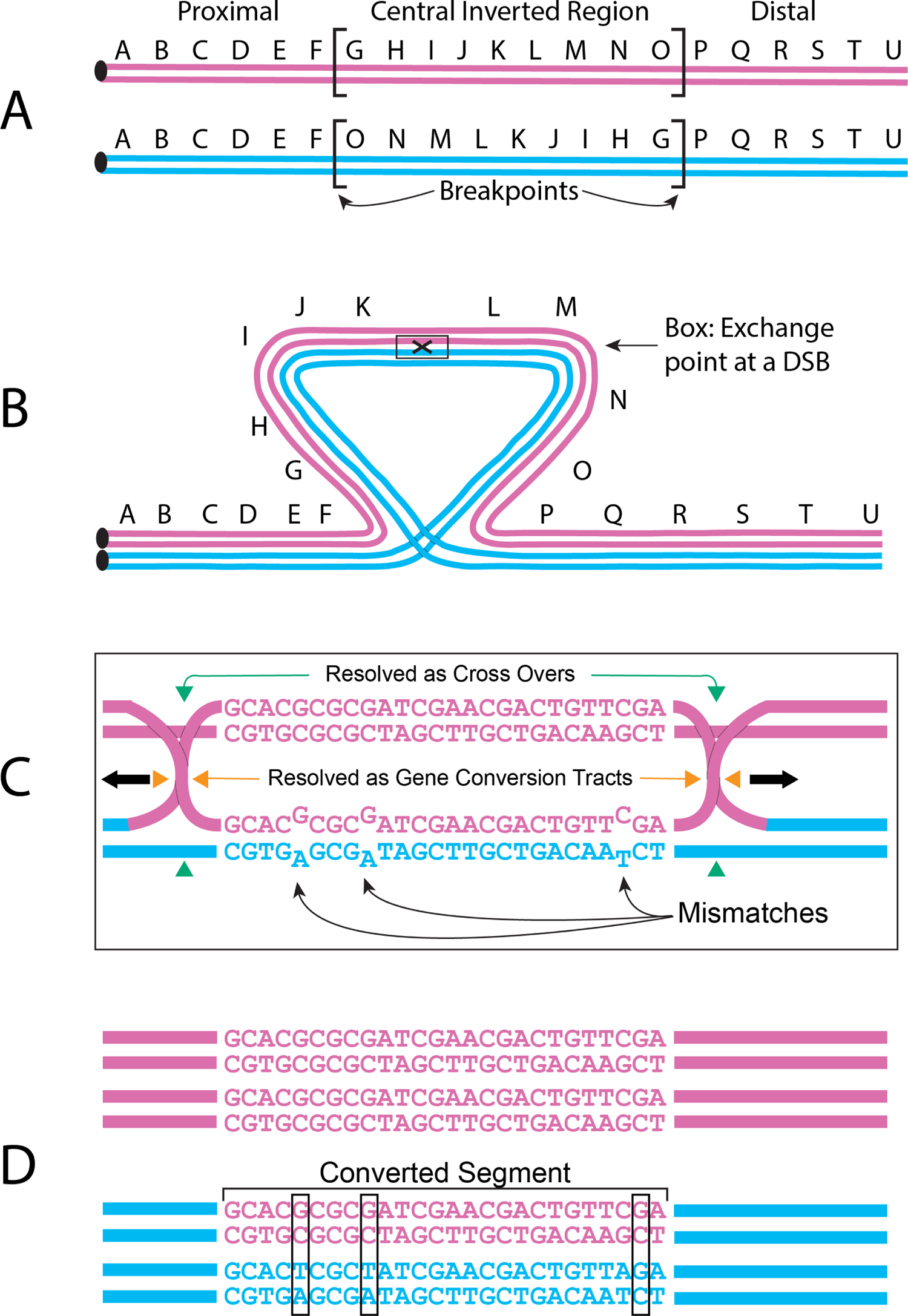
Gene conversion events generated in a heterozygote for a paracentric inversion. A. Two different gene arrangements with a paracentric inversion between the F and P genes. B. Pairing of these two inversions during meiosis showing the initiation of exchange at a double strand break (DSB within the Box). C. Exchange of strands between the two homologs via the exchange of the pink strand into the double helix of the blue strand, the so-called Holliday junction. Nucleotide mismatches are formed in this structure as DNA strands are exchanged between the homologs. The Holliday junction can be resolved either by breaking strands indicated by the green arrows leading to cross over events or by the orange arrows leading to non-cross over events and potential gene conversion tracts. D. The resolution of heteroduplex sequences because of repair. If the blue strand in the heteroduplex is used as the template for repair, then gene conversion does not occur. In this case, the pink strand was used as the template for repair leading to gene conversion event.

Fuller *et al*. (2017) analyzed complete genome sequences of the third chromosome inversion polymorphism of *D. pseudoobscura* as a model system to determine mechanisms responsible for the establishment of inversions in populations. The third chromosome or Muller C (Muller 1940) of *D. pseudoobscura* has a wealth of gene arrangements that were generated through a series of overlapping inversion events (Dobzhansky and Sturtevant 1938). These arrangements have been suggested to be targets of selection based on stable clines across heterogenous environments (Dobzhansky 1944; Anderson *et al*. 1991), stable altitudinal gradients (Dobzhansky 1948), and seasonal cycling (Dobzhansky 1948). Aquadro *et al*. (1991) used phylogenetic analysis of restriction fragment length polymorphisms to support the unique origin hypothesis of the *D. pseudoobscura* gene arrangements. Fuller *et al*. (2017) tested 2,668 Muller C genes within each of six arrangements for evidence of selection. Loci with significant elevated frequencies of derived nucleotide variants and with significantly long population specific branch lengths (Shriver *et al*. 2004; YI *et al*. 2010) were identified as outlier genes. In addition, outlier genes that were differentially expressed or had fixed inversion specific amino acids among arrangements were considered potential targets of selection.

These outlier loci may reflect regions enriched for nucleotide variation originally associated with the new inversion event that has yet to be decoupled by genetic flux. Alternatively, genetic flux in the form of gene conversion may be sufficient to homogenize genetic diversity, but purifying selection has removed deleterious conversion tracts.

Direct empirical estimates of gene conversion rates in inverted and colinear regions of *D. pseudoobscura* have been estimated to be 3.1 x 10^-5^ and 7.7 x 10^-6^ converted nucleotides per base per generation, respectively (Korunes and Noor 2019). Korunes and Noor (2019) generated genome wide SNP heterozygotes and used computational methods to infer gene conversion events on four of the five major chromosomal arms. In the cross, the third chromosome was heterozygous for the ST and PP arrangements. Ten F_2_ offspring were sequenced for this cross and 11 and 23 gene conversion events were scored within inverted and colinear regions respectively. Identical events were scored in the cross and were presumed to occur during mitosis prior to meiosis (Korunes and Noor 2019). If only unique events are scored, then three and eight gene conversion tracts were observed in inverted and colinear regions, respectively, which reduces the estimates of gene conversion rates in inverted and colinear regions to 8.5 x 10^-6^ and 2.4 x 10^-6^ converted nucleotides per base per generation, respectively. These estimates of gene conversion rates are two orders of magnitude greater than the mutation rate, which would allow gene conversion to homogenize genetic diversity among arrangements. Korunes and Noor (2019) show that gene conversion events can be detected after a single generation, but do not provide insights about the selective effects of these events, which might be inferred over longer evolutionary time.

A general computational method to detect gene conversion events indirectly was developed based on the configuration of polymorphic nucleotides (phylogenetic partitions) at multiple sites across a sample of nucleotide sequences (Stephens 1985). This method was refined to detect gene conversion events in inversion systems when differentiated DNA segments are exchanged between one arrangement and another (Betran *et al*. 1997). Figure 2 shows an example of two detected gene conversion events on Muller C of *D. pseudoobscura*. The Betran *et al*. (1997) method assumes that the frequency of an informative variant is greater in the source arrangement than in the recipient. Two probabilities are used to infer gene conversion tracts from samples of gene arrangement sequence data. The first is the probability that a SNP is informative of a conversion event based on the frequencies of a variant within the source and recipient subpopulations or ψ. The second is the probability that a gene conversion event is extended to an additional nucleotide or φ. This method was applied by Schaeffer and Anderson (2005) to gene arrangement samples of short aligned sequences (339 to 517 bp) from seven gene regions of *D. pseudoobscura*. They estimated the mean conversion tract length to be 205 bp and mean population conversion rate to be 3.4 x 10^-6^, which is comparable to the rates inferred by Korunes and Noor (2019).

**Figure 2.**
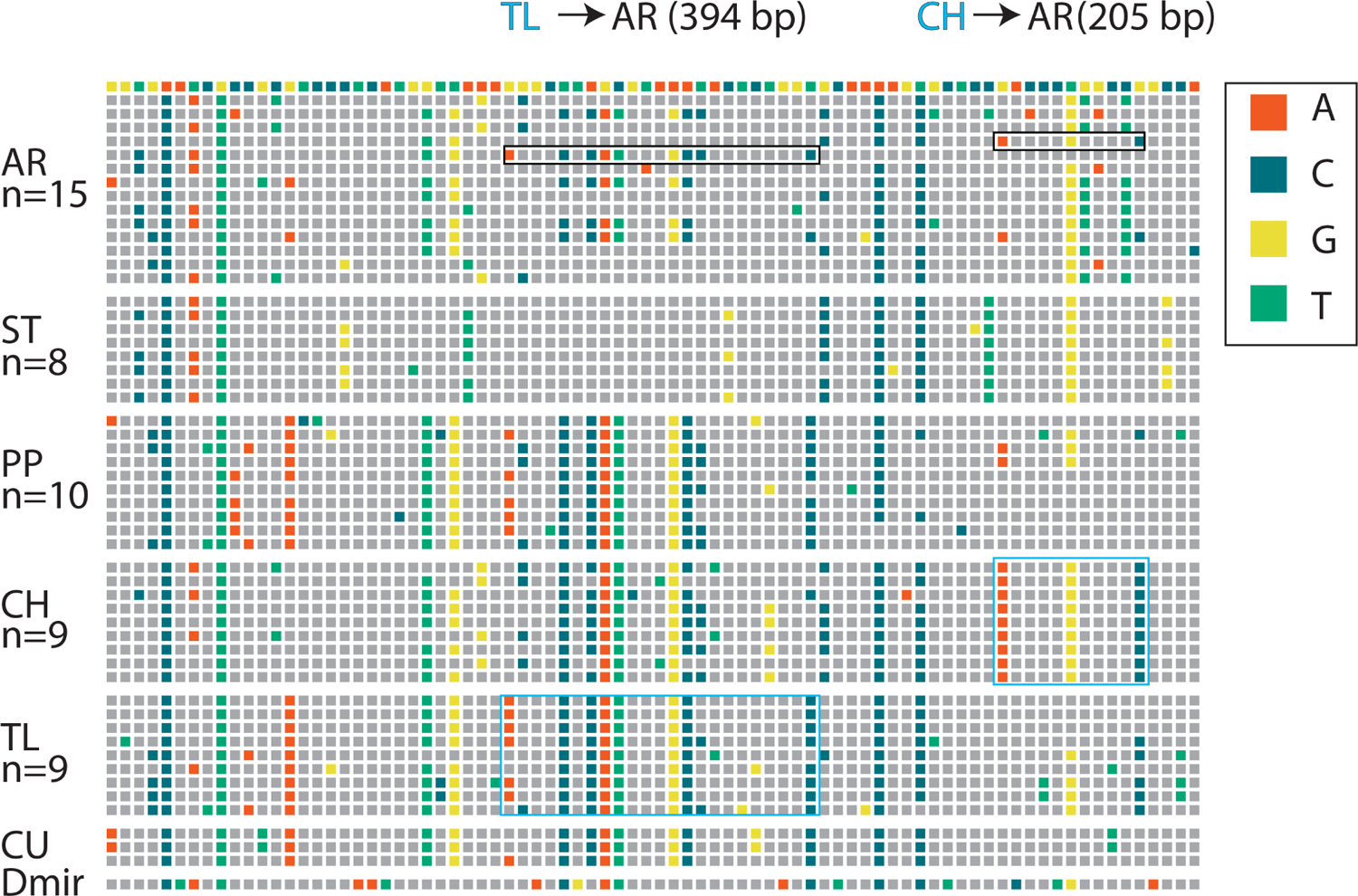
Detection of gene conversion. A map of SNP polymorphism is shown for a short region of the third chromosome of *D. pseudoobscura*. Each column represents a SNP with gray boxes matching the reference genome in the first row. A nucleotide that differs from the reference genome is represented in one of four colors (see legend on the right-hand side). Multi-site genotypes are shown for six gene arrangements along with their sample sizes: Arrowhead (AR), Standard (ST), Pikes Peak (PP), Chiricahua (CH), Tree Line (TL), Cuernavaca (CU), and the outgroup *D. miranda*. Two inferred gene conversion events are shown with the black outline box indicating the recipient sequence and the light blue outline boxes shown in light blue representing the source sequence. The length of the conversion tract is determined from the first and last nucleotide coordinate of the detected tract.

Under a neutral model and an infinite amount of time, maximal nucleotide divergence will be found at inversion breakpoints with the central inverted region showing less divergence among different arrangements (Navarro *et al*. 2000; Noor *et al*. 2007; Guerrero *et al*. 2012). When an inversion captures locally selected alleles, selection opposes the homogenizing effect of genetic flux leading to local peaks of differentiation (Guerrero *et al*. 2012). The observation of peaks of differentiation will become more apparent as time progresses and gene conversion events homogenize variation among arrangements except at selected loci.

Five *D. pseudoobscura* arrangements had sufficient sample sizes to detect multiple outlier loci. These gene arrangements had between 36 and 149 outlier genes that were distributed within and outside the boundaries of the derived inversion (Figure 1). Outlier genes had decreased estimates of the recombination parameter rho (ρ) (Chan *et al*. 2012) compared to intervening non-outlier loci (see Table 3 in Fuller *et al*. 2017) suggesting that genetic flux is limiting differentiation at a fine scale. The predominant force homogenizing local regions among arrangements, however, remains unclear. The current study infers gene conversion events over the accumulated history of the Muller C inversion polymorphism in *D. pseudoobscura* to test the hypothesis that gene conversion events occur uniformly across all genes in colinear and inverted regions of Muller C in *D. pseudoobscura*. With genome re-sequencing data, we used the Bertran *et al*. (1997) method to do a comprehensive analysis of gene conversion tracts across Muller C of *D. pseudoobscura*. We show here that gene conversion has been sufficient to homogenize genetic variation among arrangements within inverted regions. Previously identified outlier genes tended to have lower coverage from gene conversion events than non-outlier genes. These data provide evidence for the indirect effect of recombination maintaining associations of selected genes originally captured by inversion events. Outlier genes that have low gene conversion coverage represent loci where purifying selection has removed exchanged DNA.

## Materials and Methods

### Strains

The genome sequencing of 54 strains of *D. pseudoobscura* that carry one of six different gene arrangements on Muller C have been previously described (Fuller *et al*. 2017). In brief, isofemale strains were collected from seven localities: Mount St. Helena, CA (MSH), Santa Cruz Island, CA (SCI); James Reserve, CA (JR); Mather, CA (MA); Kaibab National Forest, AZ (KB); Davis Mountains, TX (DM), San Pablo Etla, Oaxaca, Mexico (SPE). Crosses with either the Blade or Lobe third chromosome balancer strains were used to generate isochromosomal strains (Drosophila Species Stock Center Strain ID 14011-0121.173 is the Blade balancer for non-AR chromosomes, Genotype: or Bl cv AR/lethal CU; Strain ID 14011-0121.171 is the Lobe balancer for non-SC chromosomes, Genotype: or L SC/ or + lethal ST; and Strain ID 14011-0121.172 is an additional strain used in the balancer crosses, Genotype or pr) (see Figure 1 in Dobzhansky and Queal 1938 for the general crossing scheme). The 54 strains used in this study carry one of six Muller C gene arrangements where the names are based on the locality where the chromosome was first collected (Dobzhansky and Sturtevant 1938): Arrowhead (AR, n=15); Standard (ST, n=8); Pikes Peak (PP, n=10); Chiricahua (CH, n=9); Tree Line (TL, n=9); and Cuernavaca (CH,n=3).

### Genome Sequences

Each of the 54 strains were sequenced with Illumina HiSeq short reads (for full details of the sequencing, read mapping, and variant calling, see Fuller *et al*. 2017). Reads were mapped to the *D. pseudoobscura* v3.2 Arrowhead reference sequence. The Arrowhead MV2-25 reference genome is available at NCBI in BioProject PRJNA18793, the re-sequencing genomes are available in BioProject PRJNA358242. Two additional Pikes Peak genomes are available at NCBI as BioSamples SAMN00709017 and SAMN00709004. The set of 54 aligned sequences for the 19.8 Mb third chromosome (Muller C) were used in the gene conversion analysis. The mapping of Illumina reads might be expected to decrease with increasing inversion distance between arrangements, however, the different inverted chromosomes have similar levels of inferred gaps or missing data (2.0 to 2.2 % sites of the 19.8 Mb sequence) and have similar distributions of gaps or missing data across the 73 regions examined in the gene conversion analysis (See Supplemental Figure S1).

### Gene Conversion Analysis

We estimated gene conversion tract coverage in outlier and non-outlier genes with the following analysis. Coverage refers to how often conversion tracts occur at a particular nucleotide in a gene region. As gene arrangements age, they accumulate arrangement specific mutations. These SNPs can be exchanged between arrangements in heterokaryotypes via crossing over or gene conversion, but will have lower frequency in the recipient strain. We used the approach of Betran *et al*. (1997) to infer gene conversion tracts across Muller C of *D. pseudoobscura*. This approach compares sets of sequences among different chromosomal arrangements to identify two or more SNPs in linkage disequilibrium that are found in two or more arrangements.

The gene conversion events detected by the Betran *et al*. (1997) approach are based on exchange between gene arrangement heterokaryotypes. The gene conversion analysis requires that the SNPs use a common gene order and that the distance between SNPs be preserved in the data set. The seven pairs of *D. pseudoobscura* inversion breakpoints used in this study divide the third chromosome into 14 regions where genes within the region are in the same order in all arrangements (Figure 3). While conversion events may occur near breakpoints (CROWN *et al*. 2018), they are unlikely to straddle the breakpoints.

**Figure 3.**
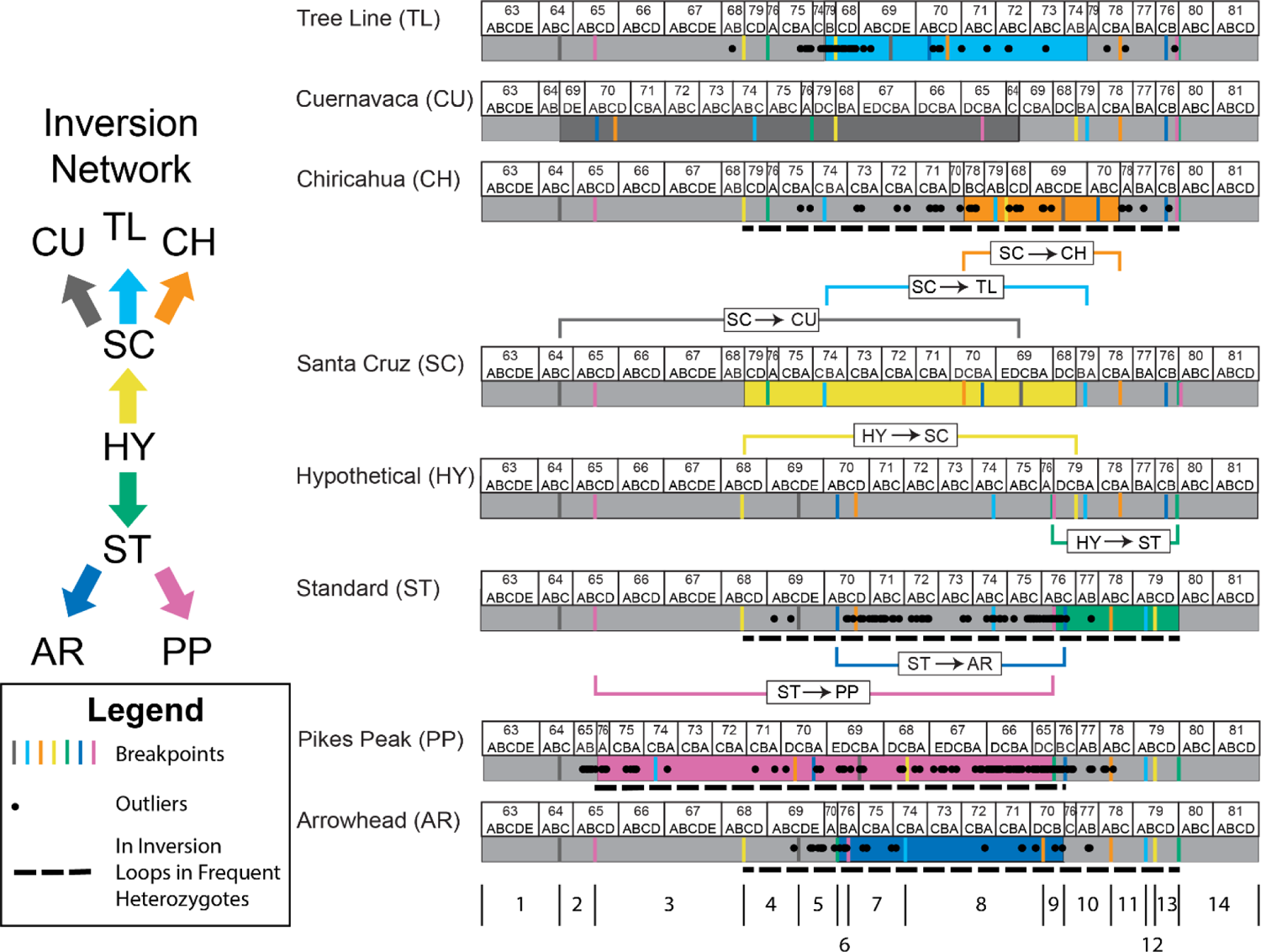
Network of Drosophila pseudoobscura gene arrangements. The network of gene arrangements (HY, ST, AR, PP, SC, CU, TL, and CH) and the inversion events (colored arrows) are shown to the left where the HY arrangement is the ancestral arrangement. One the right side are the cytogenetic maps for the gene arrangements along with the inverted segments indicated by the colored brackets. The color-coded lines below each arrangement show the locations of the different inversion breakpoints. Outlier genes in each of the gene arrangements are indicated by a black dot. An outlier gene had a significant elevation of derived alleles or population specific branch lengths and had either fixed amino acid differences or differential expression. Below the AR arrangement, 14 regions where genes are in the same order among all chromosomes in the set. The dotted line below the CH, ST, PP, and AR arrangements indicates the regions that would be inverted in ST/AR, ST/CH, AR/CH, or AR/PP heterozygotes whose frequency is at least 10% at least one locality in the Southwestern United States (Dobzhansky 1944).

The Gene Conversion analysis module within DNASP v 6 (Rozas *et al*. 2017) was used to infer gene conversion tracts across the 14 cytogenetic regions of Muller C. Each of the 14 regions were subdivided into smaller subregions of 181 to 440 kb each, with most subregions being 250 kb. This was done to accommodate DNASP memory constraints and to maximize the speed of analysis. All pairs of gene arrangements were compared in each of the 73 subregions with the Gene Conversion analysis in DNASP (15 total comparisons: AR-ST, AR-PP, AR-CH, AR-TL, AR-CU, ST-PP, ST-CH, ST-TL, ST-CU, PP-CH, PP-TL, PP-CU, CH-TL, CH-CU, and TL-CU). Each analysis excluded nucleotide sites with alignment gaps and the output provided lists of conversion tracts within strain, the beginning and end coordinates of the tract, the tract length excluding gaps, mean value of ψ, number of informative sites, and a list of sites with its associated value of ψ. The detected events are unidirectional in the pairwise analysis. The polarity of events is based on the informative sites being fixed or nearly fixed in the source arrangement and at low or intermediate frequency in the recipient arrangement. For instance, in the comparison of AR and CH, an event detected in the AR recipient strain will have CH as the source arrangement. Results of each of the 1,095 pairwise analyses were stored in separate text files and were also combined into a single file for subsequent summary analyses.

A gene conversion tract can be detected multiple times in this analysis. There are two sources of redundancy. First, a gene conversion tract can be inferred from multiple pairwise comparisons of gene arrangements. For instance, a gene conversion tract in an AR strain might be inferred from a comparison of AR with CH or with TL. Only one instance of this detected event is used. For the analysis of event polarity, only events detected with a single pairwise comparison is used so that we can unambiguously infer the source and recipient. Second, a gene conversion tract can be inferred in multiple strains of the same gene arrangement. It is assumed that gene conversion tracts result from unique events and finding the same event in multiple strains is a result of the tract increasing in frequency within the arrangement (Betran *et al*. 1997). Only one tract from a single strain is used when the conversion event is found in multiple strains. The effect of removing redundant gene conversion tracts from the data set is a conservative adjustment that lowers the overall gene conversion tract coverage per site in both outlier and non-outlier genes.

A histogram of gene conversion coverage in each of 2,668 genes was constructed from the set of non-redundant gene conversion tracts (Figure 4). Outlier and non-outlier transcripts were mapped to Muller C (Figure 4A). Next, gene conversion tracts were mapped to Muller C (Figure 4B). Finally, a histogram of gene conversion coverage was constructed by counting the number of tracts that cover each nucleotide across the chromosome (Figure 4C).

**Figure 4.**
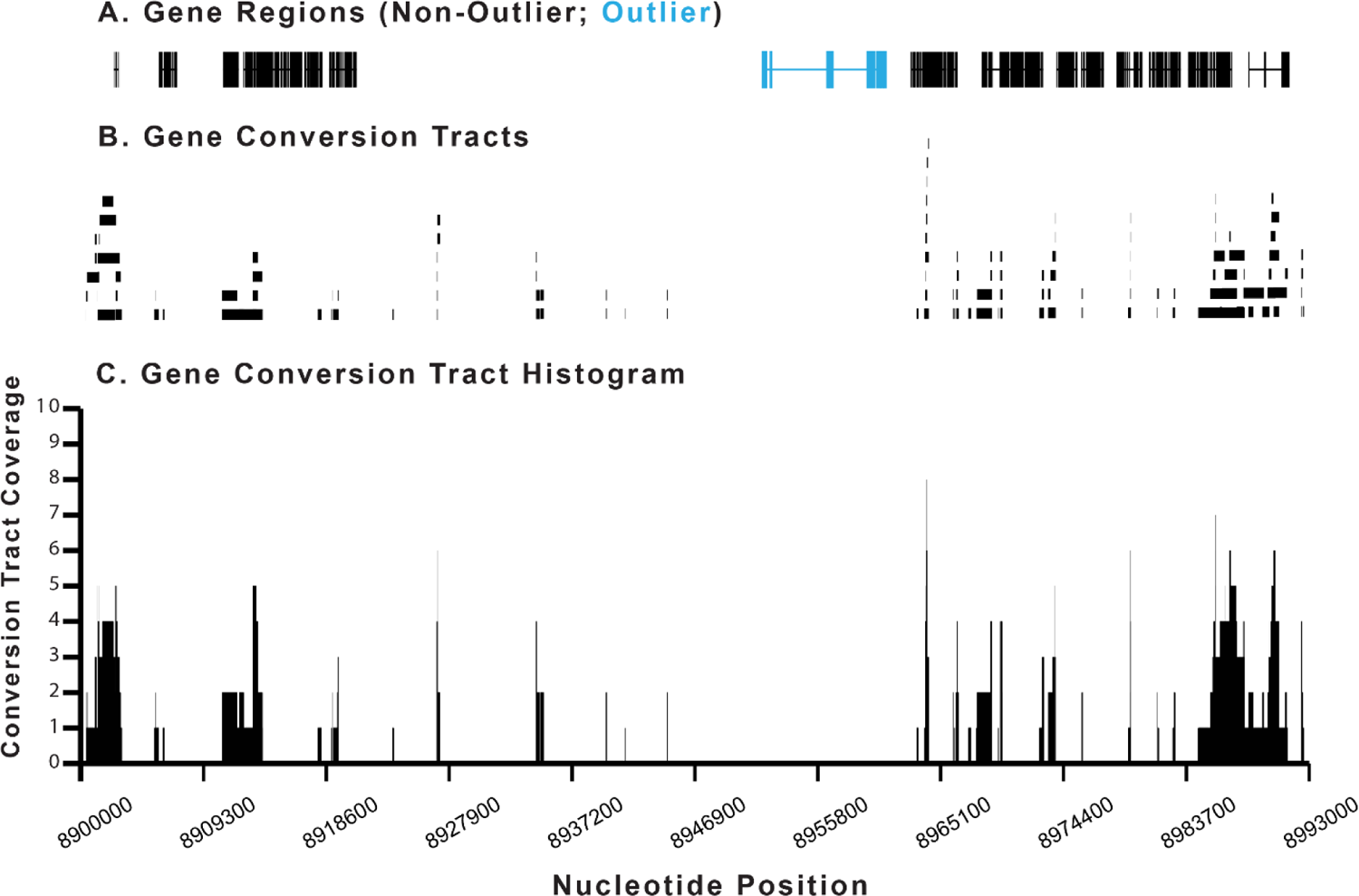
Example of a gene conversion map from the AR chromosome Coordinates 8,900,000 – 8,993,000. A. The locations of non-outlier (black) and outlier genes (blue) are shown in this section of the Muller C nucleotide sequence. B. The positions of mapped gene conversion tracts are shown on the Muller C nucleotide sequence. C. The gene conversion tracts are converted to a histogram based on the number of tracts that cover each nucleotide of the Muller C sequence.

The mean coverage per nucleotide was estimated for outlier and non-outlier genes from the gene conversion histogram. The sum of gene conversion tract coverage was estimated across the exons of each gene and divided by the total number of nucleotides in the gene. The mean gene conversion coverage was determined separately for outliers and non-outlier genes.

To determine if the donor and recipient conversion events were independent across the 14 syntenic block regions and overall, we permuted the donor and recipient classifications across the 14 regions. We constrained the permutations so that only non-self-arrangement events occurred. A total of 10,000 random permutations were used. For each permutation of the donors and recipients, the number of gene conversion events was re-estimated and compared with the observed data to determine the probability of obtaining more extreme values as well as the minimum and maximum values found across all permutations.

### Maximum Likelihood Estimates of φ and Expected Number of Conversion Tracts

We used the maximum likelihood approach of Betran *et al*. (1997) to infer φ within each of the 14 regions and for each donor-recipient pair using the observed list of conversion tracts for each pair (See equations 4, 6, and 8). The second derivative of the likelihood equation was used to estimate the asymptotic variance, which is used to infer a lower and upper bound for tract length. We did separate analyses for each region and donor-recipient pair because we assumed that the breakpoints that defined the 14 regions might interfere with the progression of conversion tracts. This may be a conservative assumption because we define the syntenic block regions based on the set of 13 breakpoints of gene arrangements used in this study.

### Tests for Differences in Gene Conversion Tract Coverage between Outlier and Non-Outlier Genes

We tested the hypothesis that outlier and non-outlier genes had equivalent gene conversion coverage using several methods. This assumes that gene conversion can occur uniformly across the chromosome.

1. *Random Permutation of Gene Class.* We used a permutation test where we randomly assigned outlier and non-outlier designations to the genes within the arrangement. A total of 10,000 replicates were used to determine the frequency of random outlier assignments that had a mean tract coverage less than the observed value.
2. *Random Permutation of Observed Gene Conversion Tracts.* We randomly assigned the observed gene conversion tracts to Muller C without replacement 10,000 times and for each random mapping we re-constructed the conversion tract histogram for the outlier and non-outlier genes. The average coverage in outlier and non-outlier genes was determined for each replicate and the frequency of replicates that had a value less than the observed was estimated.
3. *Random Gene Conversion Tracts Based on Maximum Likelihood Parameters.* We used maximum likelihood estimates of region-specific φ and the expected number of gene conversion tracts (*E(k*_T_) in the 14 different syntenic block regions to generate a set of conversion tracts. The parameter φ is the probability of extending the conversion tract to an additional nucleotide and the process of generating tract lengths is geometric random variable. For each gene conversion tract within a recipient arrangement, we sampled a tract length as a geometric random deviate. If the tract length was greater than the region size, then a new tract length was chosen. Next, a random nucleotide site was chosen from a uniform distribution to begin the gene conversion tract. If the mapped tract spanned two syntenic blocks, a new starting coordinate was chosen. A total of *E*(*k*_T_) tracts are generated for each region for each of the five donor arrangements, where *k*_T_ is the expected number of conversion tracts (See page 95 in BETRAN *et al*. 1997). A total of 10,000 replicates was generated and the mean tract coverage in outlier genes was re-estimated.

It is possible that the original classification of Muller C genes into outlier and non-outlier categories may reflect ascertainment bias because they were originally identified based on higher levels of divergence. To overcome this possibility, we classified each gene into one of three categories based on the random permutation minimum and maximum coverage estimates. The three categories are: (1) below the permuted minimum coverage; (2) between the permuted minimum and maximum coverage; and (3) above the permuted maximum coverage. We used a Chi-square test of homogeneity for each arrangement to determine if the distribution of outlier and non-outlier genes into gene coverage classes is similar.

## Results

### Numbers and Lengths of Conversion Tracts in 14 Syntenic Block Regions

There were 98,443 non-redundant gene conversion tracts observed across Muller C. The mean number of tracts per strain is 1930.2. Sample sizes varied among arrangements, which could bias the number of gene conversion tracts detected. The distribution of gene conversion tracts rejects the null hypothesis that conversion tracts are proportional to the number of strains sampled within each gene arrangement using a Chi-Square Goodness of Fit test (*X*^2^=2281.0, *df*=4, *P*<0.0001). The AR, CH, and TL arrangements have more gene conversion tracts than expected while the PP and ST arrangements have fewer tracts than expected (Figure 5). Gene conversion tract numbers are not similar in the 14 syntenic block regions in the five different arrangements based on a Chi-Square test for Homogeneity (*X*^2^=366.6, *df*=52, *P*<0.0001) (Figure 6). Examination of residuals shows that the AR arrangement has a deficiency of tracts in regions 5 and 9, the CH arrangement has an excess of gene conversion tracts in region 5 and a deficiency of tracts in regions 11 and 14, the PP arrangement has a deficiency of tracts in region 5 and an excess of tracts in region 9, the ST arrangement has deficiencies of tracts in regions 3 and 5 and an excess of tracts in regions 8 and 10, while the TL arrangement has an excess of tracts in region 3 and deficiencies of tracts in regions 5 and 8.

**Figure 5.**
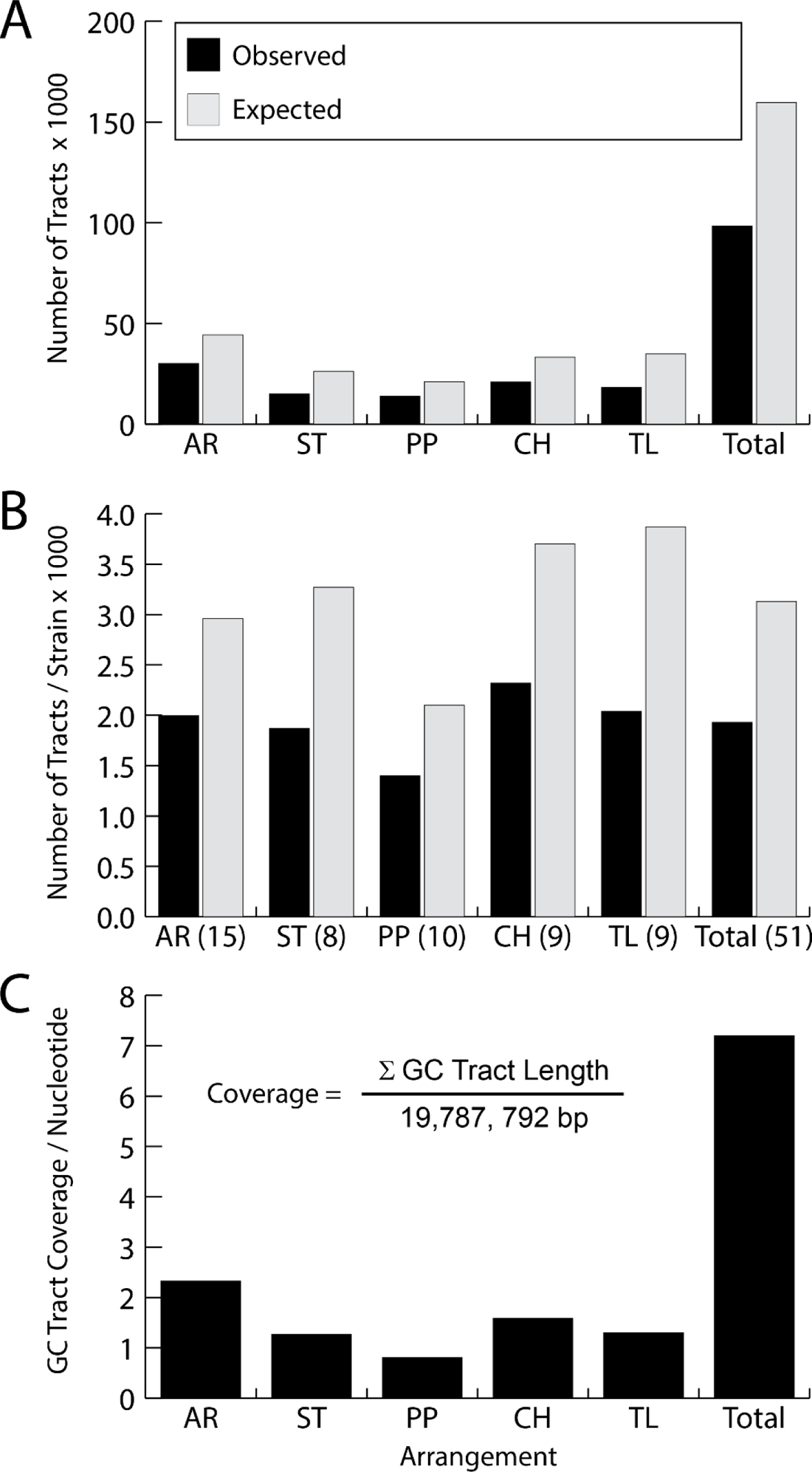
Gene conversion tract information in five Muller C gene arrangements in D. pseudoobscura. A. Observed and expected numbers of gene conversion tracts where the expected number = Number of Observed/[1-Probability(Unobserved Gene Conserved Events)]. B. Observed and expected mean number of gene conversion tracts per strain where the number of strains is shown in parentheses next to the gene arrangement name on the x-axis. C. Gene conversion tract coverage per nucleotide for the five gene arrangements.

**Figure 6.**
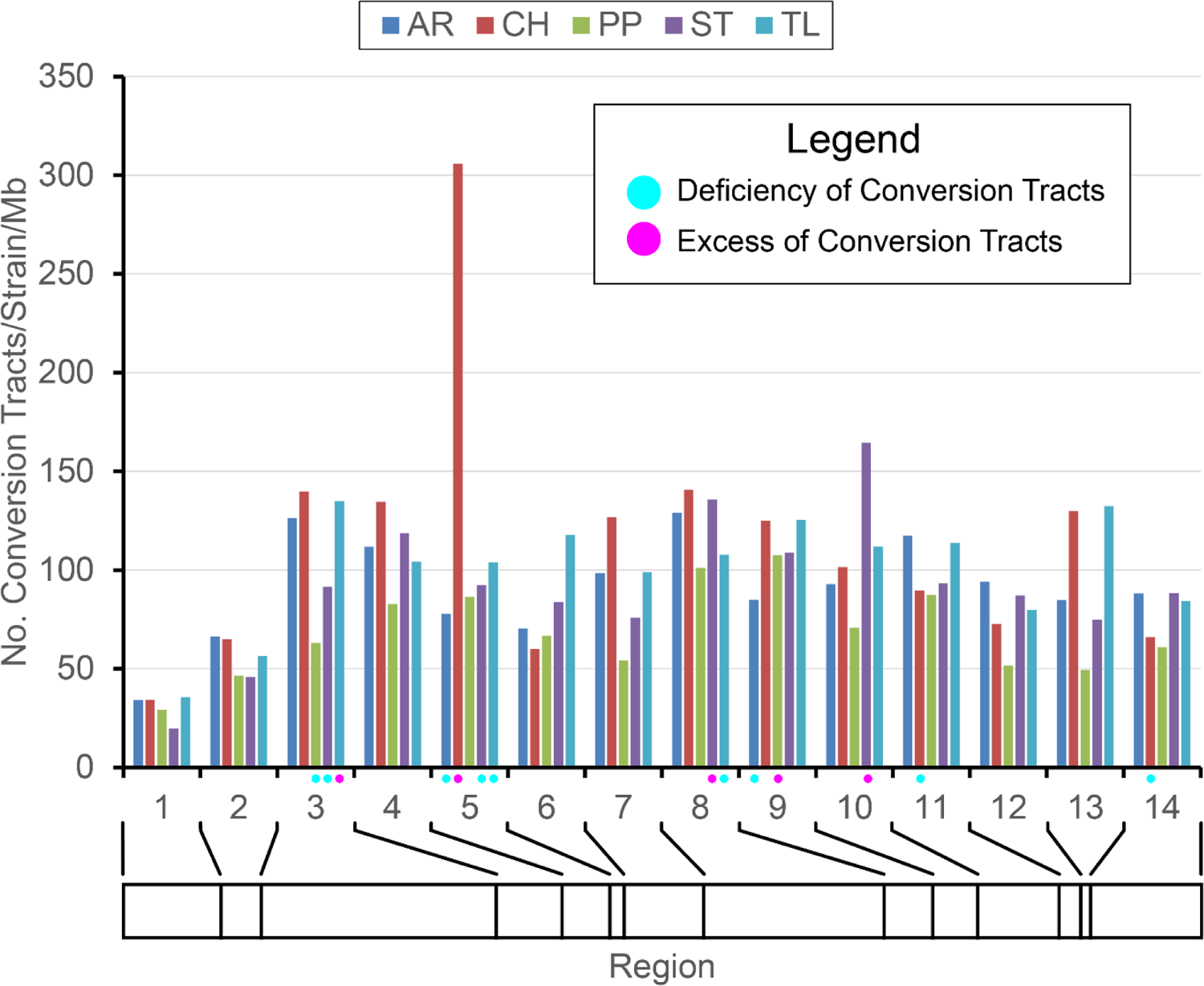
Distribution of the number of conversion tracts within recipient gene arrangement for 14 regions along Muller C of D. pseudoobscura. Dots underneath the x-axis indicate regions within gene arrangement with a deficiency (blue) or excess of gene conversion tracts based on residuals from a Chi Square test of homogeneity. The bar underneath the x-axis shows the relative sizes of the different regions.

Median observed gene conversion tract lengths are greatest in the proximal and distal regions of Muller C compared to the central inverted regions (Figure 7). The central areas of the chromosome which are often within inverted regions in heterokaryotypes have an observed median tract length of 112, 160, 97, 127, and 115 bps in the AR, CH. PP, ST, and TL arrangements, respectively. We used a random permutation test to determine if the median gene conversion tract lengths are lower or greater than expected given the distribution of all observed tract lengths. The test was performed by shuffling the list of tracts without replacement, which randomly assigns tracts to the 14 different regions keeping the number of tracts per region the same as the observed. For each of the 10,000 permutation replicates, the regional median was re-estimated and the probability of obtaining a median greater than or less than the observed value within each region was determined. Figure 5 shows the regions that had median gene conversion tract lengths that were significantly less than or greater than expected given the distribution of all tract lengths. Proximal regions 1 and 2 and distal regions 12 through 14 had median tract lengths that were greater than expected in two or more gene arrangements while central regions 4 through 10 had median tract lengths that were less than expected in four or more karyotypes. Regions 3 and 11 had arrangements with median tract lengths less than or greater than expected reflecting boundary regions that the transition between largely collinear or inverted segments of the chromosome.

**Figure 7.**
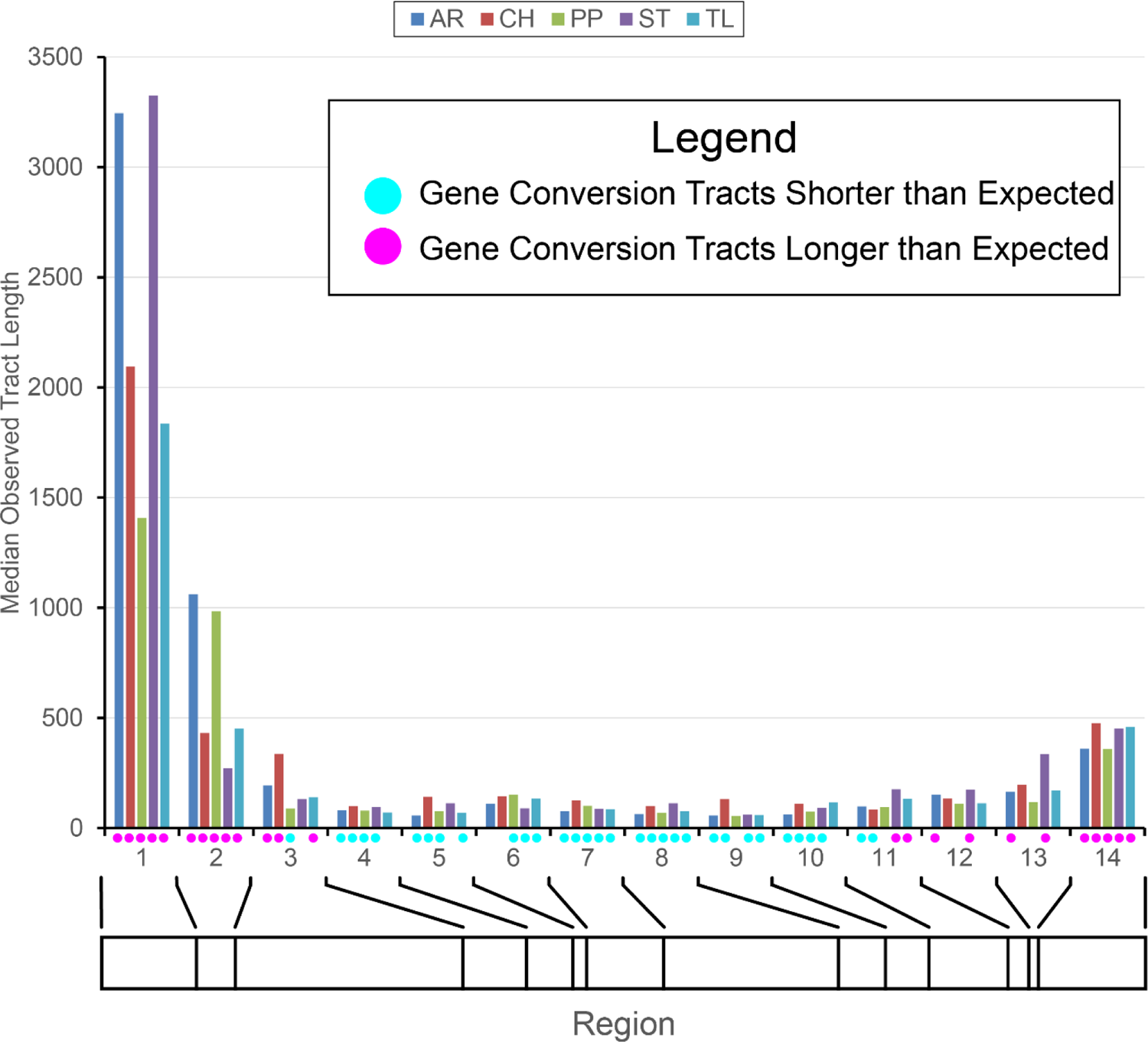
Distribution of conversion tract lengths within recipient gene arrangement for 14 regions along Muller C of *D. pseudoobscura*. Dots underneath the x-axis indicate regions within gene arrangement where gene conversion tract lengths are shorter (blue) or longer (pink) than expected based on a random permutation test. The bar underneath the x-axis shows the relative sizes of the different regions.

Of the 98,443 observed gene conversion tracts in recipient arrangements, the source arrangement could be inferred unambiguously in 74,419 events. We observed gene conversion events between all pairs of arrangements across Muller C ranging between 1,270 conversion events from ST to PP and 5,266 conversion events from CH to AR. Figure 8 shows the polarity of the observed gene conversion events (Donor to Recipient) in the 14 syntenic block regions and across all regions. For all gene conversion events across all regions, three pairs showed a deficiency of events in both directions (AR-TL, CH-PP, and ST-TL), one pair showed an excess of events in both directions (AR-ST), one pair showed a deficiency in one direction and an excess in the other direction (PP-ST).. The rest showed either a deficiency or excess in only one direction. The test of independence of donor to recipient events for the 14 different regions found subsets of regions with significant deficiencies or excesses of conversion events with a tendency toward what was found for the sum of all events. For instance, if the overall trend was toward a deficiency of events, then most regions with a significant departure tended to also show a deficiency, e.g., the ST to CH transition had a significant overall deficiency (2,195) and ten of 14 regions showed a significant deficiency. For the arrangement pairs likely to form heterozygotes at frequencies of 10% or more in natural populations (CH/ST, CH/PP, AR/ST, and AR/PP) CH tended to be a donor to AR and ST, ST tended to be a donor to AR, and PP tended to be a donor to AR.

**Figure 8.**
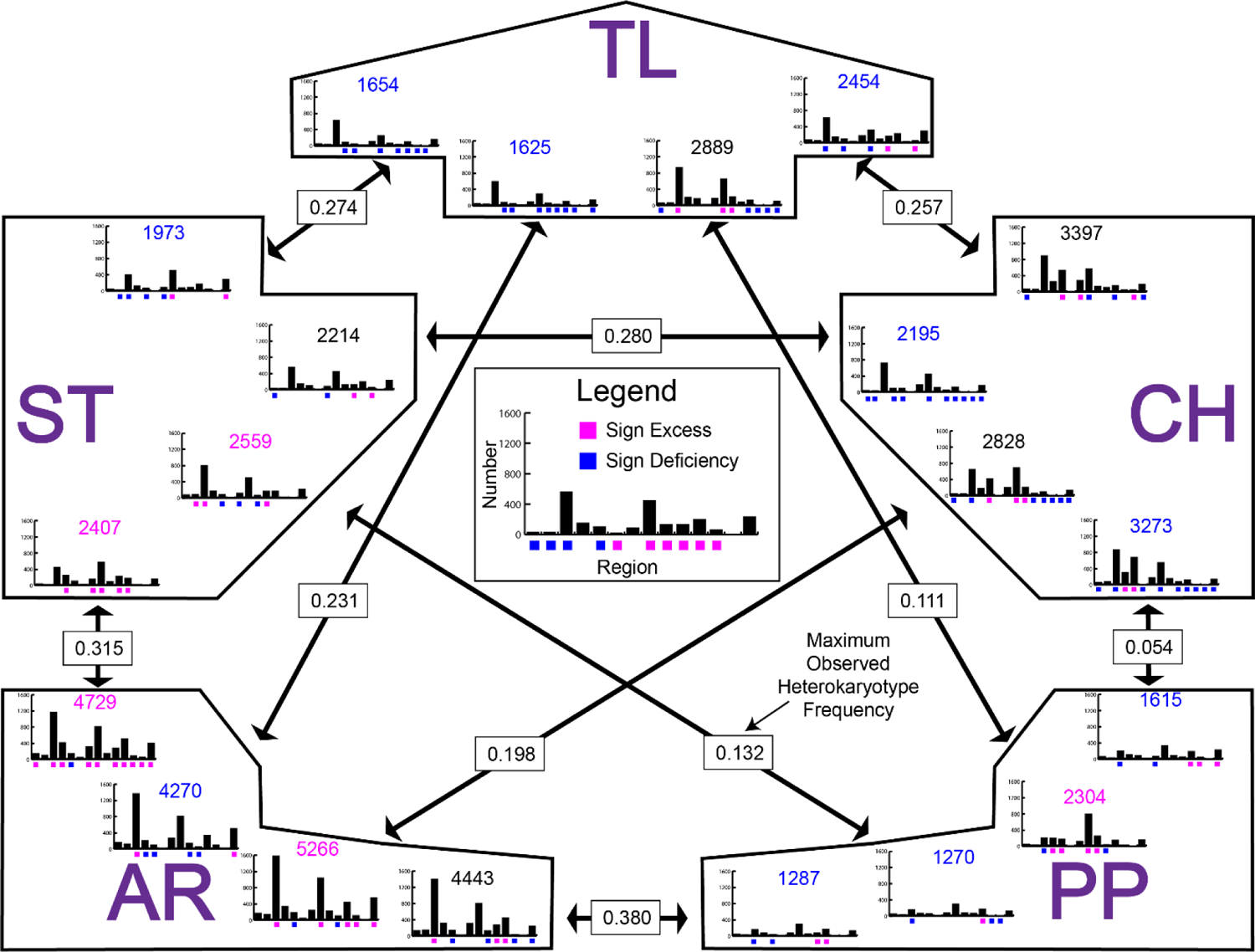
Number of gene conversion events per strain between pairs of gene arrangements in *D. pseudoobscura* in 14 different syntenic blocks. The five gene arrangements are abbreviated with their two letter codes: AR, Arrowhead; CH, Chiricahua; PP, Pikes Peak; ST, Standard; TL, Tree Line. Histograms show the number of gene conversion events in the 14 syntenic blocks with the maximum value on all scales being 2,500. Blue and pink boxes below each bar indicate regions with a significant deficiency (blue) or excess (pink) of events based on a random permutation test (see the text). The number above each histogram indicates the total number of events across all 14 regions with a blue number indicating a significant deficiency, a pink number indicating a significant excess of events, and a black number indicating neither a significant deficiency or excess. The double headed arrows connect pairs of arrangements with the arrows pointing to the number of events in the recipient arrangement from the donor arrangement. For instance, 1654 conversion events were from ST to TL and 1973 conversion events were from TL to ST. The boxes with frequencies on the arrows are the maximum observed heterokaryotype frequencies in the 1980 collections of *D. pseudoobscura* (Anderson *et al*. 1991).

### Maximum Likelihood Estimates of φ and Expected Number of Conversion Tracts

The estimates of φ and the expected number of conversion tracts along with other values for all donor-recipient pairs in each region are in the Supplemental Materials (Supplemental Table S1). These values are used below in tests of gene conversion coverage in outlier genes.

### Gene Conversion Tract Coverage in Outlier and Non-Outlier Genes

The 98,443 observed conversion tracts were mapped to Muller C and the mean coverage per nucleotide was estimated for the exons of each transcript. The expected gene conversion tract coverage per nucleotide across Muller C is 1.2 per nucleotide for all arrangements except for PP, which had an expected coverage rate of 0.8 tracts per nucleotide. Thus, at least one conversion tract should occur in each gene across the chromosome (Figure 5).

Figure 9 shows the gene conversion tract coverage for outlier and non-outlier genes across Muller C in five gene arrangements of *D. pseudoobscura*. Proximal and distal regions of all arrangements have higher conversion tract coverage than central inverted regions. Outlier genes tend to have lower levels of coverage than non-outliers. The maximum coverage values are sample size dependent. Standard, Chiricahua, Pikes Peak, and Tree Line have sample sizes between 8 and 10 with maximum coverage of 10 tracts/nucleotide. Arrowhead, on the other hand, has a sample size of 15 with a maximum coverage of 18.

**Figure 9.**
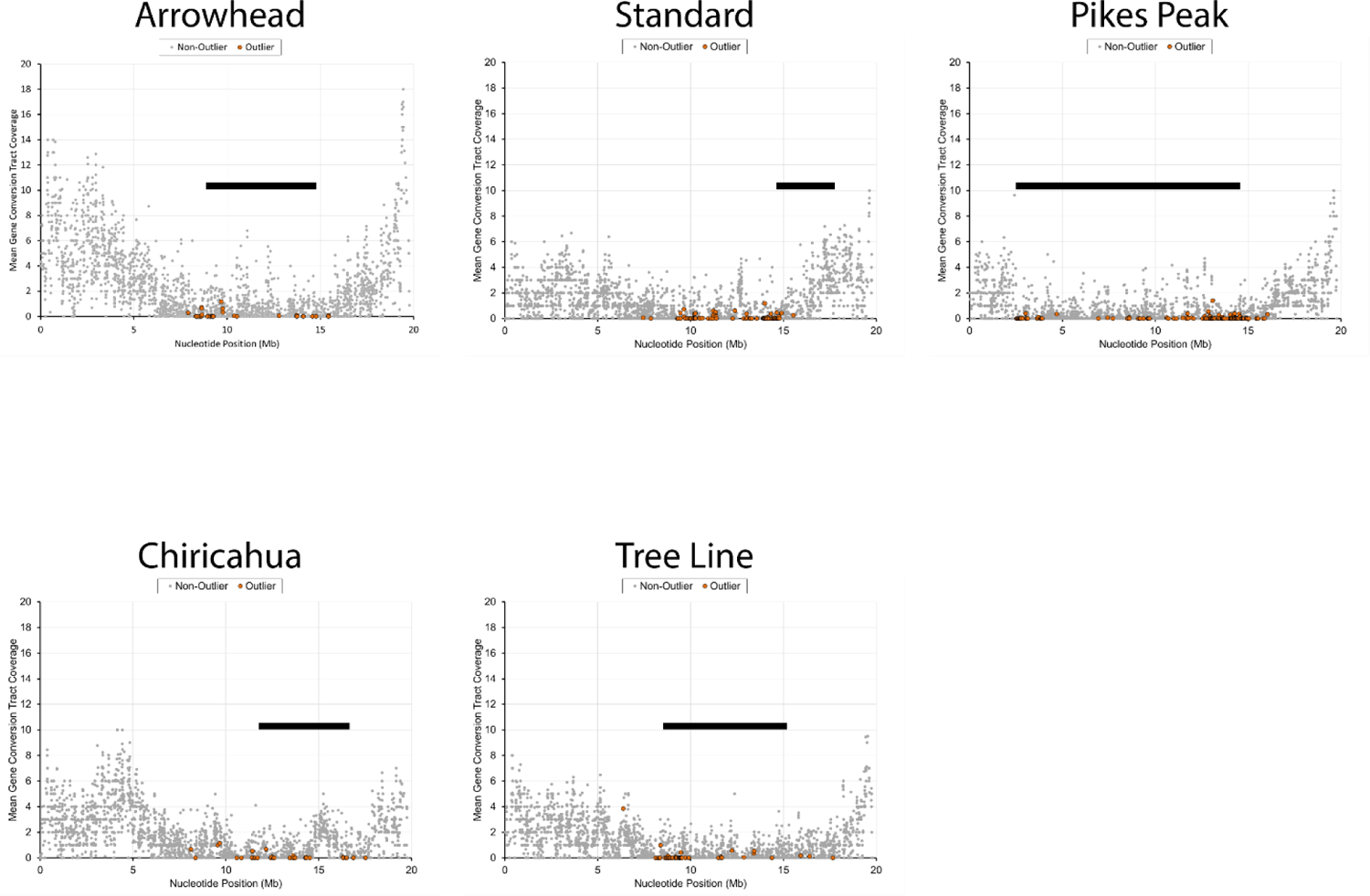
Mean gene conversion tract coverage in 2,668 transcripts across Muller C in *D. pseudoobscura* in five gene arrangements. Coverage is the number of times that a nucleotide is covered by a gene conversion tract. Coverage was estimated for each exon within a transcript and the mean coverage for the entire transcript was averaged over all exons. Outlier genes are shown with an orange marker while non-outliers are shown with a gray marker. The location of the derived inversion giving rise to each gene arrangement are shown with a black bar.

The mean gene conversion tract coverage estimated for the 2,668 genes on Muller C shows that non-outlier genes have higher mean coverage than outlier genes (Table 1). Gene conversion tract coverage in non-outlier genes matches the expected value for the entire chromosome, while outlier genes have coverage values < 0.1.

**Table 1.** Gene Conversion Tract Coverage of Outlier and Non-Outlier Genes on Muller C of *Drosophila pseudoobscura*.

| Gene Arrangement | Outlier<br>Obs Coverage (SD) | Non-Outlier<br>Obs Coverage (SD) | Min – Max |
| --- | --- | --- | --- |
| Arrowhead (AR) |  |  |  |
| Transcripts | 36 | 2,632 |  |
| Nucleotides | 78,468 bp | 4,099,525 bp |  |
| Permute Out | 0.112 (0.503)* | 2.585 (3.064) | 1.406 – 4.037 |
| Shuffle CT |  |  | 0.679 – 2.652 |
| ML Parameter |  |  | 2.018 – 4.066 |
| Standard (ST) |  |  |  |
| Transcripts | 82 | 2,596 |  |
| Nucleotides | 179,724 bp | 3,998,269 bp |  |
| Permute Out | 0.067 (0.356)* | 1.415 (1.662) | 0.974 – 1.926 |
| Shuffle CT |  |  | 0.563 – 1.758 |
| ML Parameter |  |  | 1.415 – 2.565 |
| Pikes Peak (PP) |  |  |  |
| Transcripts | 149 | 2,519 |  |
| Nucleotides | 365,249 bp | 3,812,744 bp |  |
| Permute Out | 0.048 (0.295)* | 0.922 (1.488) | 0.609 – 1.156 |
| Shuffle CT |  |  | 0.233 – 0.922 |
| ML Parameter |  |  | 0.922 – 2.242 |
| Chiricahua (CH) |  |  |  |
| Transcripts | 40 | 2,628 |  |
| Nucleotides | 95,262 bp | 4,082,731 bp |  |
| Permute Out | 0.055 (0.322)* | 1.735 (1.873) | 1.087 – 2.709 |
| Shuffle CT |  |  | 0.483 – 1.735 |
| ML Parameter |  |  | 1.595 – 3.199 |
| Tree Line (TL) |  |  |  |
| Transcripts | 54 | 2,614 |  |
| Nucleotides | 113,586 bp | 4,064,407 bp |  |
| Permute Out | 0.101 (0.444)* | 1.290 (1.656) | 0.832 – 1.917 |
| Shuffle CT |  |  | 0.216 – 1.290 |
| ML Parameter |  |  | 0.658 – 1.660 |

\*, $P < 0.0001$ the observed coverage value for the outlier genes is less than that observed in the randomly permuted data. Min – Max, minimum and maximum values from 10,000 replicates; Permute Out, permutation test where outlier label was permuted; Shuffle CT, permutation test where the observed conversion tracts were shuffled; ML Parameter, test where the expected number of conversion tracts with random tract length with maximum likelihood parameter $\phi$ is used.

We used three tests to determine if the lower gene conversion coverage is expected by chance (Table 1). The Random Permutation of Gene Class test found that the observed mean gene conversion coverage in outlier genes was always less than random permutations in all gene arrangements. One factor to consider is that we filtered the conversion tracts to remove redundant tracts. If the redundant tracts were to be included, we would likely see higher coverage both in outlier and non-outlier genes, but still show a significant difference. The Random Permutation of Observed Gene Conversion Tracts test showed that the observed mean coverage in outliers was always less than the replicate permutations. The Random Gene Conversion Tracts Based on Maximum Likelihood Parameters test found that the observed mean conversion tract coverage in outlier genes was less than was found with random conversion tracts. The results of this test should be viewed with caution because the expected number of true tracts is greater than the observed number of tracts. Thus, it is not surprising that we observe higher gene conversion tract coverage in outlier genes than in our replicates. The important result from these three tests is that outlier genes have lower gene conversion coverage than expected.

It is possible that the previous classification of genes into outlier and non-outliers may bias our analysis of gene conversion tract coverage because these genes were identified based on high levels of fixed nucleotide mutations, which would have low levels of shared polymorphism that would result from gene conversion. We examined the relationship between the Population Specific Branch Length (PSBL) statistic and gene conversion tract coverage in outlier and non-outlier genes in the five gene arrangements (Supplement Figure S2). Non-outlier genes had gene conversion tract coverage estimates from 0 to 18 while non-outlier genes had tract coverage estimates from 0 to 4. The random permutation analysis provides an overall estimate for the minimum and maximum coverage across all genes. We divided genes up into three classes, (1) below the permuted minimum value, (2) between the permuted minimum and maximum value, and (3) above the permuted maximum value. We did this separately for the outlier and non-outlier genes and used a chi-square test of homogeneity to determine if the distribution into the three classes is the same for outlier and non-outlier genes (See Supplemental Figure S2 A-E and Tables S2 to S6). The Chi-square tests show that the distributions reject homogeneity in the five arrangements (Supplemental Tables S2-S6). For the five gene arrangements, the outlier class has an excess of genes with gene conversion tract coverage below the permutation minimum (Percent AR = 94.4%; CH=87.5%; PP=92.6%; ST=95.1%; TL=88.9%). Each arrangement had classified outlier genes with coverages between the permuted minimum and maximum coverages (Numbers, AR, 2; CH, 5; PP, 10; ST, 4; TL, 5) and two arrangements had classified outlier genes above the permuted maximum coverage (Numbers, PP, 1; TL, 1), which suggest that some previously identified outliers do experience expected levels of gene conversion.

### Gene Conversion and the Site Frequency Spectrum

We examined whether the site frequency spectrum differs in regions with different levels of gene conversion. Gene conversion events would likely introduce maladaptive genetic variants and our analysis suggests that these variants are removed by purifying selection, however, purifying selection does not have as profound effect on the site frequency spectrum as beneficial mutations do. Beneficial mutations will sweep out nucleotide variation likely leading to an excess of rare variants and a significant negative Tajima’s D. Purifying selection on deleterious variants will only remove or decrease the frequency of chromosomes that carry the changes. Jackson et al. (2015) found that purifying selection alters the site frequency spectrum of nonsynonymous nucleotides, but did not alter the site frequency spectrum of synonymous sites. Supplemental Figure S3 A-E shows volcano plots of Tajima’s D for all genes of the five arrangements with significant and non-significant D indicated in pink or blue for outlier genes. The five arrangements have both significant positive and negative Tajima’s *D*s with only the CH having more positive than negative D values.

## Discussion

The work presented here shows that gene conversion events among arrangements are readily detectable and serve as a source of genetic flux among the gene arrangements of *D. pseudoobscura*. The number of observed conversion tracts tends to be uniformly distributed across Muller C (Figure 6) except for the proximal two syntenic blocks of the chromosome consistent with theory on genetic flux (NAVARRO *et al*. 1997). Fewer conversion tracts may result from the initiation of fewer double stranded breaks in the proximal regions. While regions 1 and 2 are closer to the centromere where crossing over is reduced (LINDSLEY AND SANDLER 1977), these regions are in euchromatic DNA, which is unlikely to be affected by the centromeric effect. More likely is that the Betran *et al*. (1997) method has lower power to detect gene conversion events in regions 1 and 2. This method relies on informative nucleotide polymorphisms to detect an exchange between two gene arrangements. Because regions 1 and 2 are outside most inverted regions, genetic exchange is likely to reduce differentiation and prevent conversion tract detection.

The observed median gene conversion tract lengths are largest in the proximal and distal regions of Muller C and are reduced in size in the inverted regions (Figure 7). The detected events in the proximal and distal regions may be the result of cross overs rather than gene conversion tracts. The Betran *et al*. (1997) method uses informative nucleotide sites to infer exchanges between arrangements. The accumulation of genetic differences among gene arrangements can extend beyond the boundaries of the inversion breakpoints providing information about gene exchanges. In this case, the exchange events are longer than what is observed within the inverted regions. The most proximal region (1) has the largest median conversion track length compared to the most distal region (14). This may reflect that region 1 is further from the influence of interference caused by inversion breakpoints (GONG *et al*. 2005). Most common arrangement pairs are homosequential across regions 1 to 3 with the exception of the PP arrangement, which has its proximal breakpoint at the boundary of regions 2 and 3.

As a result, there is likely to be less interference of inversion breakpoints in the more proximal regions leading to resolution of exchanges as cross overs and longer sequences being exchanged.

We expected exchanges among all pairs of gene arrangements because heterokaryotypes for these pairs have frequencies between 0.054 and 0.380 in at least one population of *D. pseudoobscura*. The number of gene conversion events are not positively correlated with heterokaryotypic frequency. We find larger numbers of events between arrangements with lower observed heterokaryotypic frequencies than higher heterokaryotypic frequencies.(Anderson *et al*. 1991). A total of 4,888 tracts were observed between CH and PP, which has an observed maximum heterokaryotypic frequency of 0.054 (Anderson *et al*. 1991), while ST and CH had 4409 observed tracts with an observed heterokaryotypic frequency of 0.280. We also found that in 15 of 20 cases, the number of events from donor to recipients are either lower or greater than expected in a permutation test (Figure 8). For most pairs of arrangements, there is a bias in which arrangement is the donor versus the recipient. For instance, AR is the recipient of more tracts from CH than expected while there is neither a deficiency nor excess of events in the reverse direction. (see heterokaryotype frequencies in Figure 8). The CH and TL arrangements have two and three cases of deficiencies of being gene conversion recipients. The AR, PP, and ST arrangements have both deficiencies and excesses of being gene conversion recipients. The significant deficiencies may reflect selection against exchanged segments of DNA from one background to another especially in regions within inverted regions (Figure 8).

Arrangements that receive significant excesses of gene conversion events may reflect positively selected changes. For example, the PP arrangement received an excess of events from the TL arrangement especially in regions 8 and 9, which are cytological regions 74B-70D and 70C-70B (Figure 2 and Figure 8). These two regions are unusual because phylogenetic analysis of SNPs of the 14 regions of Muller C shows that PP and TL share a more recent common ancestor than either does with their respective cytogenetic ancestor (ST for PP and SC for TL) (see Figure 4 in Fuller *et al*. 2017). This suggests that gene conversion may have introduced sequences from the TL to the PP arrangement that were important for the proliferation of the PP arrangement. Further examination of the exact nature of how the TL donated conversion tracts alter genes in the PP arrangement are needed to understand if introduced variation is the target of selection.

The rate of gene conversion is sufficient to homogenize the sequences among the gene arrangements, yet one observes differentiation of some genes (outliers) and not others across the inverted segments. Our analysis of gene conversion coverage tracts shows that gene conversion occurs less frequently in outlier versus non-outlier genes. Tests that randomly shuffle the gene conversion tracts show that there are enough events to homogenize the sequences. Each gene arrangement has fewer than 12% of outlier genes without a significant reduction in gene conversion events (see Supplemental Tables S2-S6). The reason we can detect these outlier loci is that we observe differentiation in the face of the homogenizing effect of gene flux, which works in a similar manner as gene flow. This requires that the various gene arrangements accumulate genetic differences and that there is sufficient time for gene conversion events to accumulate. The Muller C inversion polymorphism in *D. pseudoobscura* is estimated to be at least 1 million years old (Aquadro *et al*. 1991; Wallace *et al*. 2011) with the AR arrangement being the newest arrangement of the set originating 350,000 years ago. Despite being a young arrangement, our sample of AR chromosomes has over 30 thousand gene conversion events with a mean coverage of 2.33 events per nucleotide and 36 detected outlier genes with a mean coverage of 0.11. A similar pattern of reduced coverage in outlier genes is observed in the other four older arrangements.

Essentially what is observed here is an example of survivorship bias. Gene conversion events survive over time in genes within the inverted regions that can tolerate additional genetic diversity, while gene conversion events in selected outlier genes from different arrangements do not survive because they damage the important features of selected alleles within the inversion haplotype. We assume that these outlier genes are in fact genes essential for the particular gene arrangement’s adaptive phenotype. Thus, the lack of homogenization of alleles by gene conversion allows one to detect a signal of selected genes within gene arrangements despite cross over suppression in heterokaryotypes.

Each of the major gene arrangements examined in *D. pseudoobscura* have evidence for multiple outlier genes that show arrangement specific variation. We show here that the arrangement specific variation in outlier genes is observed despite the homogenizing effect of extensive genetic flux among arrangements driven by gene conversion events through the history of these chromosomes. These data are consistent with a model where the inversions captured sets of genes involved in local adaptation (Kirkpatrick and Barton 2006). We propose that the local populations of *D. pseudoobscura* in different physiographic provinces (Dobzhansky 1944; Lobeck 1948) are selected for different locally adapted phenotypes encoded by multiple genes within the arrangements. These polygenic traits were generated through shuffling of allelic variation to generate adaptive combinations in different environments. Adaptive combinations in one environment may not be beneficial in another habitat and are likely to generate maladaptive types through recombination. This creates a setting for chromosomal inversions to establish if they capture combinations of selected genes (Charlesworth and Charlesworth 1973). The set of selected genes is not likely to be recognized initially because there will be extensive linkage disequilibrium of SNPs across the inverted region from the ancestral arrangement. As the ancestral and derived arrangements diverge in sequence, conditions to recognize selected loci emerge. Gene conversion begins to homogenize the unselected genes within the inversion and with sufficient time differentiated selected genes can be recognized.

If genes within the inverted region are neutral, then we would expect to see the highest levels of genetic differentiation at or near the inversion breakpoints and not within the central regions of the inversion (Navarro *et al*. 2000; Munte *et al*. 2005; Noor *et al*. 2007; Guerrero and Kirkpatrick 2014). We observe outlier genes distributed across the inverted regions, which are not restricted to the breakpoint regions even in the largest inversion event considered in this study, Pikes Peak. The inversion that gave rise to the PP arrangement accounts for 60% of Muller C. Double cross overs would be expected to homogenize the central region of the inverted region, yet 129 outlier loci are distributed across the rearranged segment. This differentiation pattern is the so-called suspension bridge pattern where the towers of the bridge are the regions of high differentiation or outlier genes and the cables are the regions of reduced differentiation caused by the accumulation of gene conversion events.

An alternative model to the capture model is one where new inversions gain beneficial alleles within the arrangement over time. This model does not seem plausible because in this scenario, one would predict that the repeated selected sweeps that add multiple adaptive alleles would continue to remove segregating variation from the inverted region making it impossible to identify outlier genes. A majority of nucleotide sites within the arrangement would be in near absolute linkage disequilibrium at all times due to the repeated sweeps (Fuller *et al*. 2019).

In summary, we show here that gene conversion is a powerful force for genetic flux among different gene arrangements that can allow population geneticists to identify selected loci within inversions. Selected genes are discovered as differentiated loci surrounded by regions of relative homogeneity among arrangements. This approach will not work for recent inversions because sufficient time must pass for differentiation among arrangements and for gene conversion to homogenize non-selected genes. The inversion polymorphism of *D. pseudoobscura* is sufficiently old that we have identified multiple loci within each arrangement are identified as selective targets. We think this supports the model that each arrangement captured sets of locally adapted alleles. Locally adapted inversions were favored because they prevented the formation of maladaptive combinations from other locally adapted migrant chromosomes (Kirkpatrick and Barton 2006). When an adaptive QTL is discovered to map within an inversion, researchers often have a look of despair on their faces. This study suggests there is hope to identify an adaptive QTL within the confines of an inversion. The identification of selected genes within inversions is an important step in understanding the molecular basis for what led to the establishment of inversions in populations. In this case, the outlier genes within the *D. pseudoobscura* inversions fall into two broad categories, perception and detoxification genes (Fuller *et al*. 2017). Mapping gene conversion events provides an important tool for discriminating selected from non-selected genes within inverted regions of the genome.

## Acknowledgements

The data presented in this work was generated through support from the National Institutes of Health R01 GM 098478 to (S.W.S.). We thank the sequencing team at the Baylor College of Medicine Human Genome Sequencing Center for their help in sequencing data generation. The funders had no role in study design, data collection, data analysis, the decision to publish, or preparation of the manuscript.

