## Supplemental Figures S1-S3 and Tables S2-S6 for "Genomics of Natural Populations: Gene Conversion Events Reveal Selected Genes within the Inversions of *Drosophila pseudoobscura*"

Figure S1

Gap or Missing Base Colls

| Region | AR | ST | PP | CH | TL | CU |
| --- | --- | --- | --- | --- | --- | --- |
| R01a | 33126 | 33128 | 33134 | 33098 | 33133 | 33057 |
| R01b | 27544 | 27525 | 27575 | 27529 | 27575 | 27488 |
| R01c | 21218 | 21169 | 21154 | 21129 | 21182 | 21120 |
| R01d | 364 | 329 | 341 | 380 | 410 | 295 |
| R01e | 21036 | 20936 | 21048 | 21075 | 21024 | 20809 |
| R01f | 768 | 828 | 871 | 867 | 786 | 671 |
| R01g | 31520 | 31451 | 31554 | 31516 | 31484 | 31270 |
| R02a | 17956 | 17921 | 18031 | 17953 | 18008 | 17908 |
| R02b | 1074 | 932 | 1038 | 1014 | 1056 | 898 |
| R02c | 336 | 255 | 405 | 307 | 307 | 163 |
| R03a | 498 | 446 | 1007 | 520 | 387 | 193 |
| R03b | 2033 | 1886 | 2390 | 1885 | 1973 | 1818 |
| R03c | 2261 | 1955 | 3281 | 1940 | 2655 | 1654 |
| R03d | 601 | 290 | 747 | 320 | 482 | 221 |
| R03e | 1856 | 1588 | 1958 | 1589 | 1691 | 1427 |
| R03f | 532 | 404 | 840 | 476 | 583 | 322 |
| R03g | 379 | 310 | 385 | 205 | 299 | 185 |
| R03h | 396 | 275 | 385 | 316 | 318 | 103 |
| R03i | 619 | 435 | 614 | 363 | 493 | 177 |
| R03j | 3988 | 3758 | 4108 | 3664 | 3924 | 3365 |
| R03k | 4945 | 4440 | 5170 | 4476 | 4705 | 4161 |
| R03l | 1111 | 1039 | 1165 | 944 | 1087 | 854 |
| R03m | 1716 | 1417 | 1971 | 1514 | 1838 | 1244 |
| R03n | 973 | 569 | 1258 | 602 | 1079 | 381 |
| R03o | 263 | 485 | 634 | 470 | 585 | 293 |
| R03p | 562 | 634 | 1066 | 960 | 1468 | 871 |
| R03q | 17717 | 17619 | 18072 | 18133 | 18402 | 17831 |
| R04a | 545 | 460 | 893 | 837 | 1075 | 670 |
| R04b | 573 | 359 | 949 | 1041 | 1386 | 815 |
| R04c | 457 | 275 | 576 | 526 | 613 | 358 |
| R04d | 8770 | 8696 | 9114 | 9090 | 9242 | 8909 |
| R04e | 611 | 332 | 810 | 853 | 1222 | 754 |
| R05a | 6882 | 6685 | 7040 | 7045 | 7329 | 6895 |
| R05b | 354 | 327 | 791 | 923 | 1108 | 648 |
| R05c | 501 | 536 | 939 | 955 | 1478 | 665 |
| R06a | 258 | 299 | 516 | 388 | 529 | 348 |
| R07a | 1496 | 1442 | 1954 | 1858 | 2252 | 1798 |
| R07b | 3105 | 3167 | 3236 | 3232 | 3433 | 3152 |
| R07c | 7333 | 7210 | 7535 | 7341 | 7703 | 7146 |
| R07d | 2130 | 1991 | 2366 | 2299 | 2617 | 2099 |
| R07e | 3718 | 3470 | 3796 | 3896 | 4293 | 3696 |
| R08a | 2845 | 2755 | 3320 | 3098 | 3375 | 2770 |
| R08b | 8938 | 8732 | 8946 | 8897 | 9126 | 8730 |
| R08c | 376 | 552 | 1173 | 906 | 1279 | 628 |
| R08d | 630 | 465 | 691 | 779 | 916 | 437 |
| R08e | 1755 | 1646 | 2335 | 1977 | 2492 | 1637 |
| R08f | 1174 | 1109 | 1368 | 1401 | 1473 | 949 |
| R08g | 2022 | 1924 | 2220 | 2296 | 2381 | 2412 |
| R08h | 25684 | 25368 | 25705 | 25565 | 25838 | 25422 |
| R08i | 10547 | 10719 | 10768 | 10601 | 10783 | 10580 |
| R08j | 2902 | 2991 | 3093 | 2940 | 3236 | 2877 |
| R08k | 493 | 398 | 1064 | 770 | 968 | 513 |
| R08l | 4097 | 3827 | 4316 | 4053 | 4501 | 4127 |
| R09a | 391 | 434 | 1235 | 638 | 1210 | 563 |
| R09b | 835 | 777 | 1365 | 940 | 1154 | 725 |
| R09c | 1844 | 2039 | 3110 | 2895 | 3456 | 2220 |
| R10a | 1929 | 1884 | 2480 | 2243 | 2597 | 2173 |
| R10b | 2584 | 2492 | 3554 | 3633 | 4387 | 3030 |
| R11a | 1459 | 1286 | 1621 | 1655 | 1909 | 1453 |
| R11b | 555 | 426 | 923 | 583 | 899 | 574 |
| R11c | 215 | 108 | 332 | 167 | 238 | 163 |
| R11d | 1669 | 1062 | 1951 | 1232 | 1879 | 958 |
| R11e | 51063 | 50511 | 51397 | 51070 | 51701 | 50810 |
| R12a | 333 | 213 | 357 | 330 | 647 | 256 |
| R13a | 320 | 244 | 369 | 432 | 707 | 350 |
| R14a | 1523 | 1061 | 2027 | 1170 | 1873 | 909 |
| R14b | 39619 | 39243 | 40079 | 39450 | 39680 | 39066 |
| R14c | 1326 | 1175 | 1294 | 1307 | 1293 | 1215 |
| R14d | 529 | 315 | 591 | 505 | 690 | 262 |
| R14e | 2295 | 2004 | 2449 | 2064 | 2249 | 1649 |
| R14f | 240 | 130 | 165 | 162 | 241 | 86 |
| R14g | 1713 | 1690 | 1734 | 1701 | 1686 | 1673 |
| R14h | 3968 | 3943 | 4046 | 3947 | 3985 | 3867 |

Figure S2

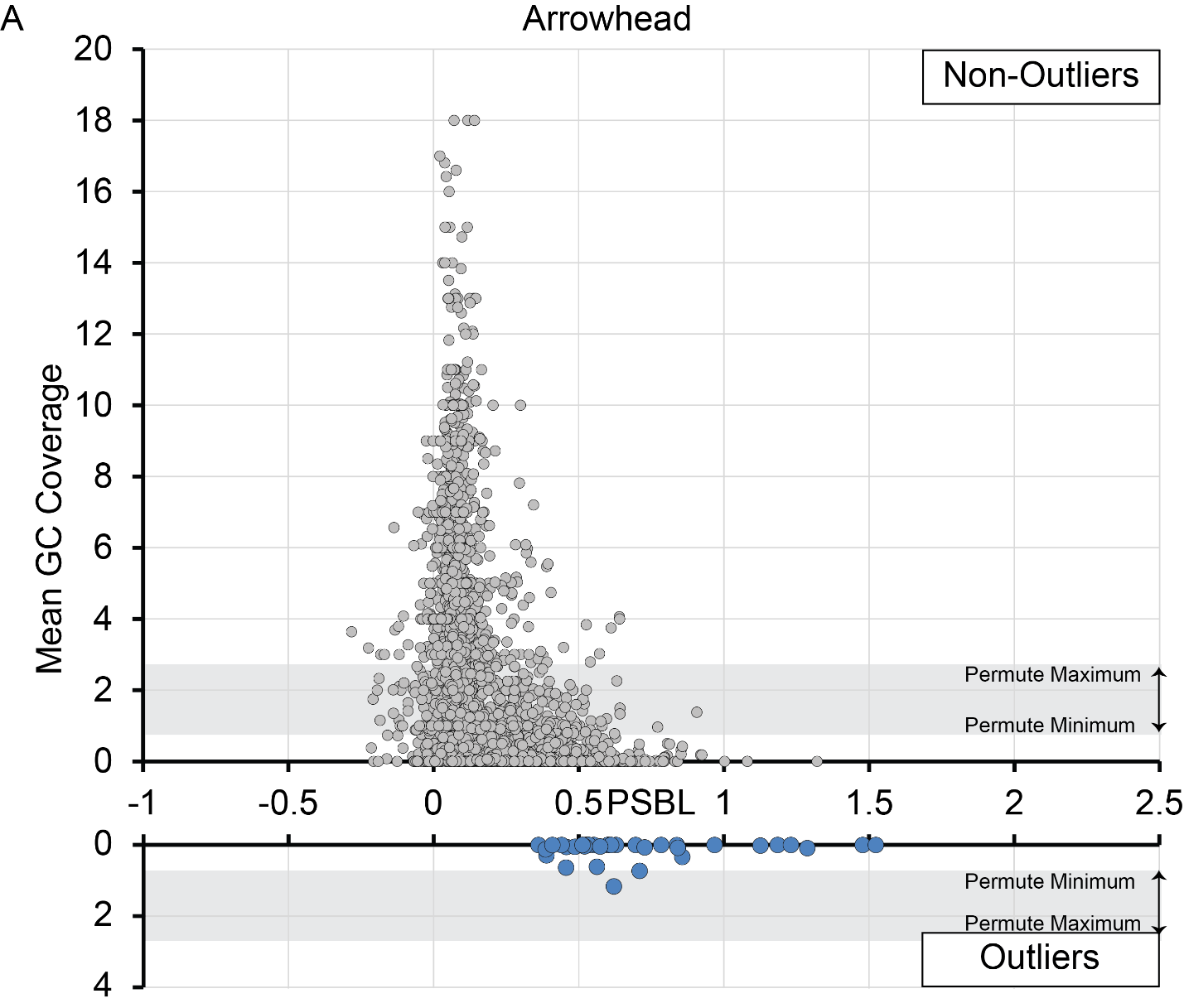

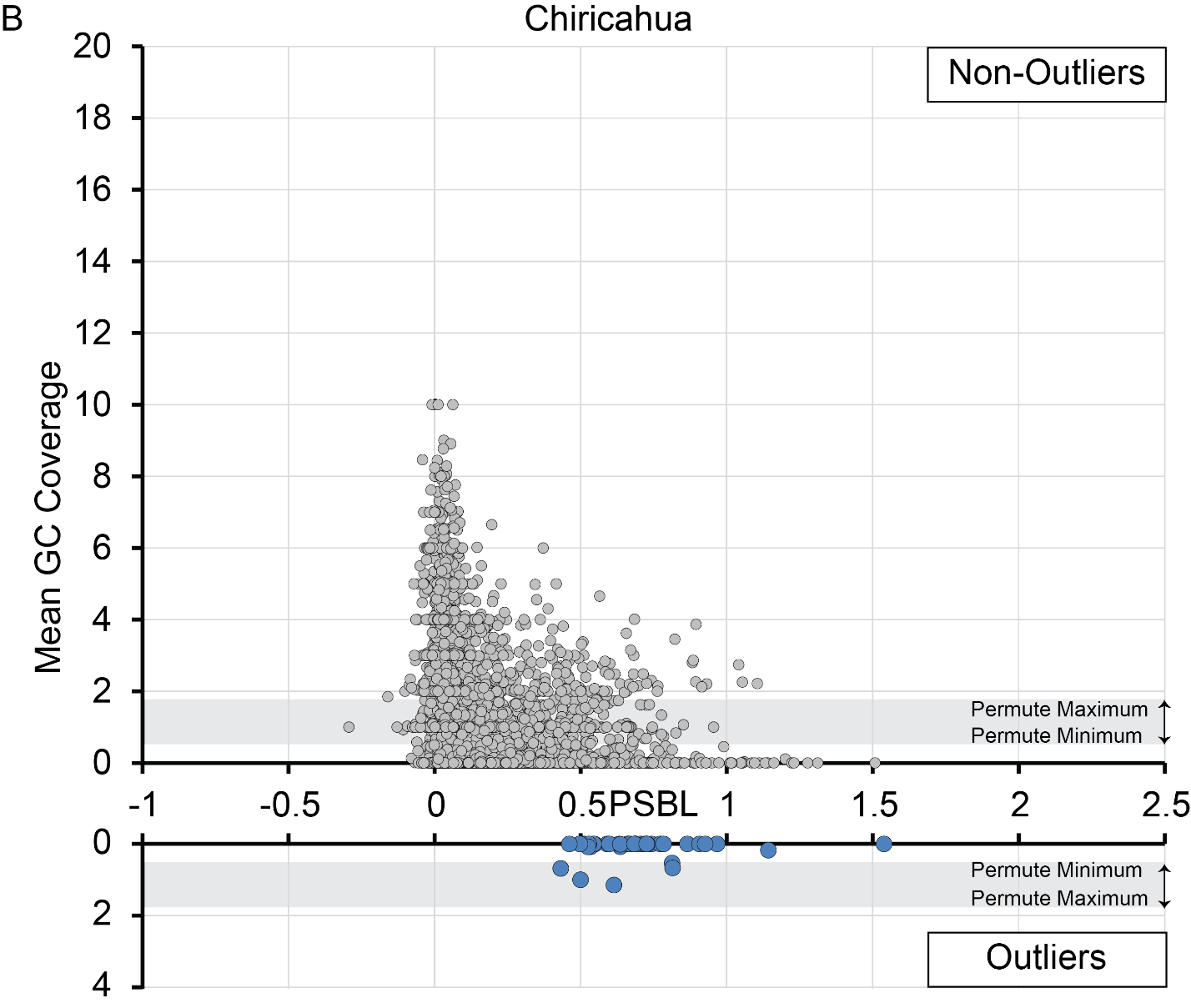

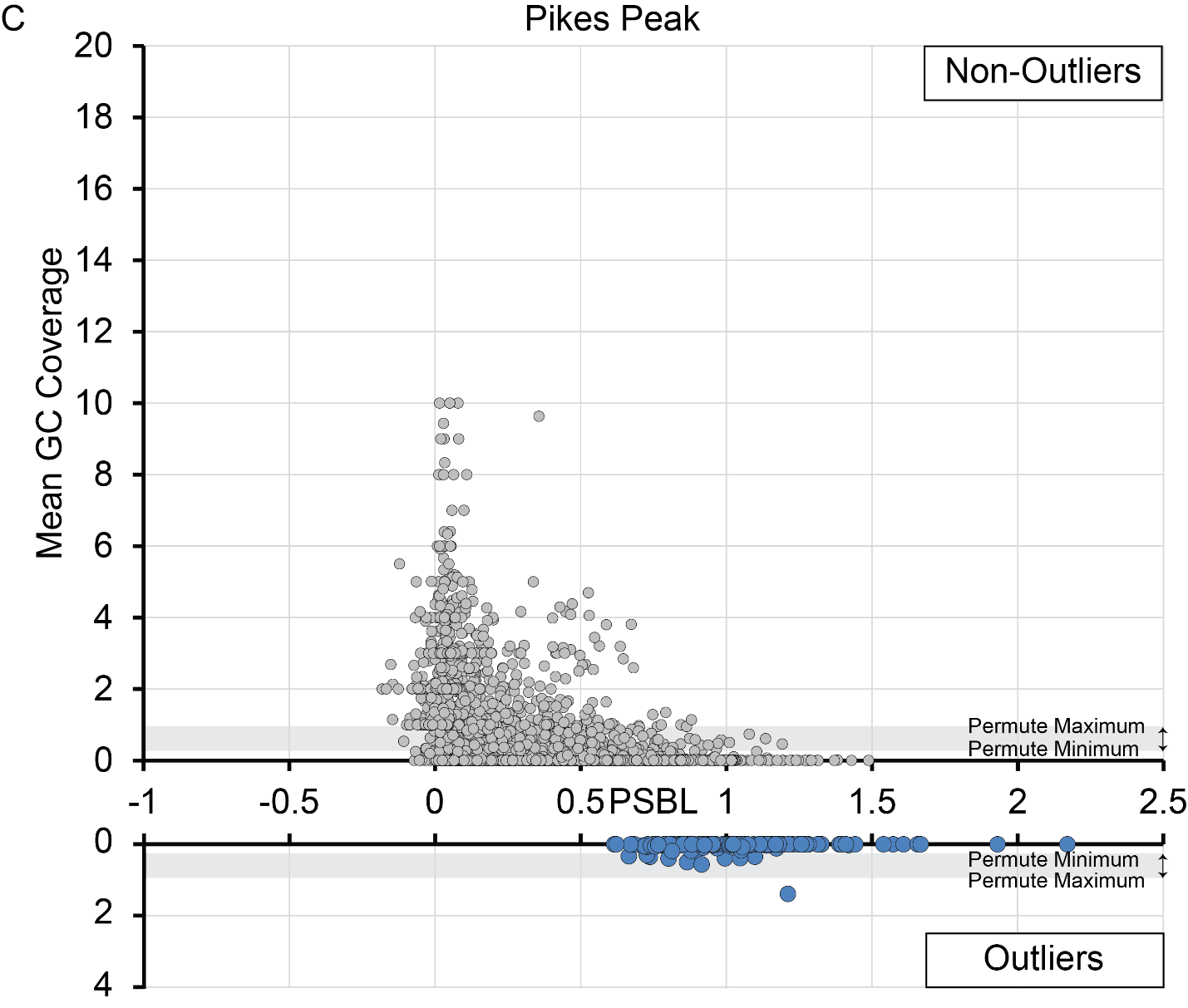

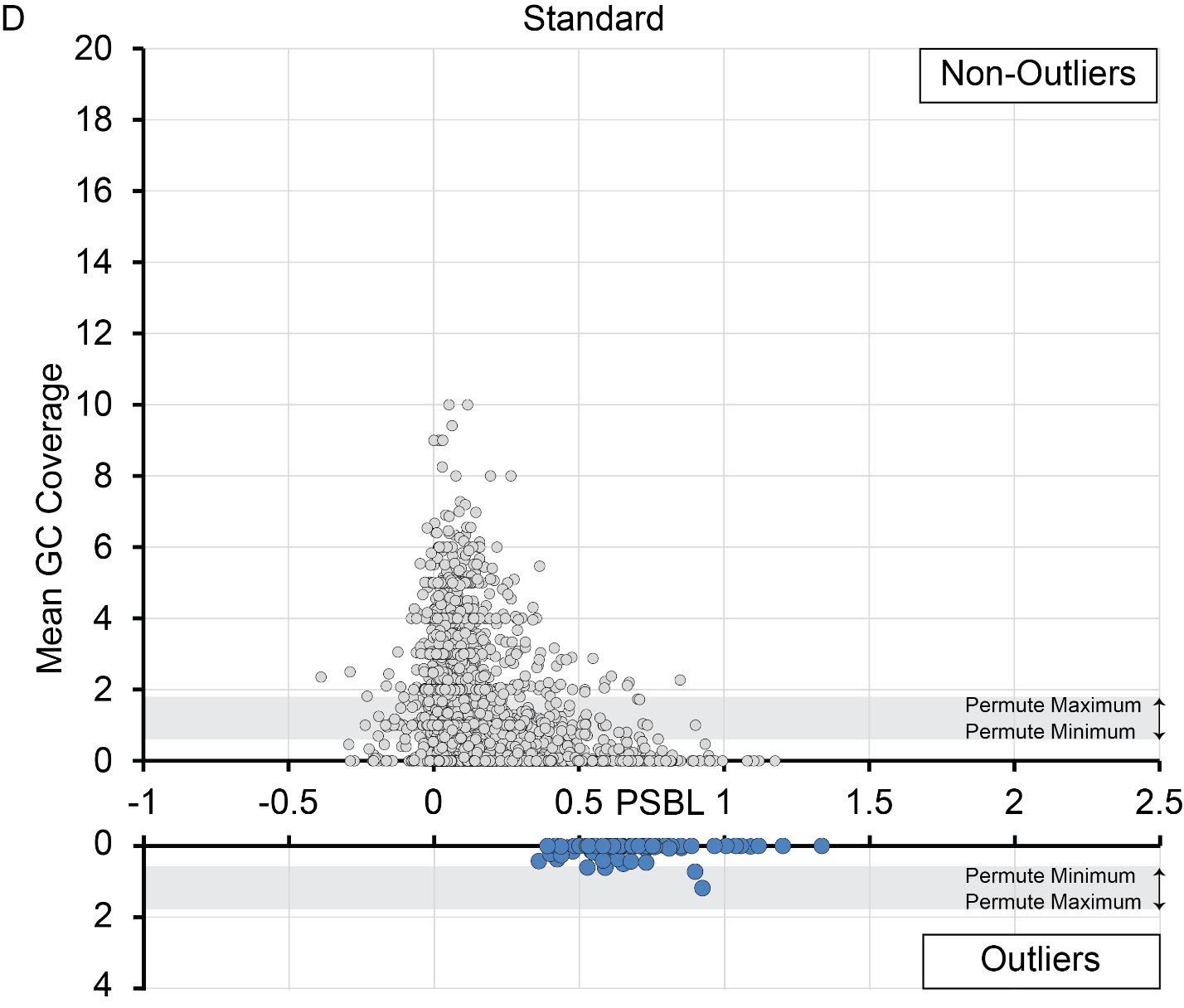

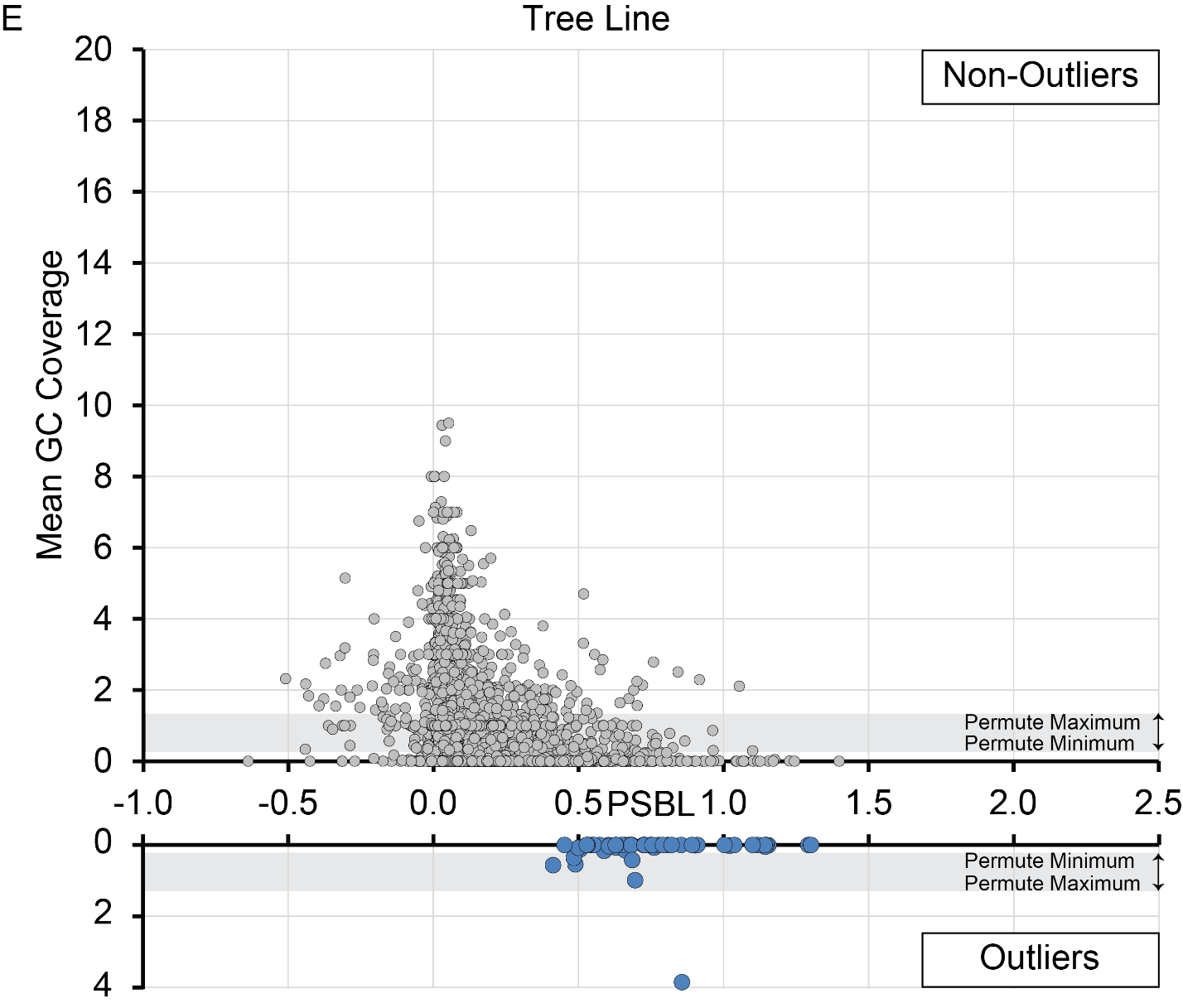

Figure S2. Relationship between Population Specific Branch Length (PSBL) on the x-axis and mean gene conversion tract coverage in 2,668 genes on Muller C of *D. pseudoobscura*. A significantly large PSBL is a proxy for a signature of selection and defines outlier genes if the gene is differentially expressed, has a fixed amino acid difference from other arrangements, or both shown as blue dots on the lower graph or gray dots on the upper graph. The gray bar highlights the minimum to maximum gene conversion coverage for genes when the gene conversion tracts within an arrangement were randomly permuted.

**Figure S3A**

Arrowhead

-Log10(P)

| Tajima's D | All Data |
| --- | --- |
| 0.28 | 3.34679 |
| 0.15 | 3.3098 |
| -1.075 | 3.08619 |
| 0.086 | 2.86967 |
| -1.68 | 2.83268 |
| 0.151 | 2.79048 |
| -1.356 | 2.44733 |
| -1.565 | 1.89894 |
| -1.031 | 1.55878 |
| -0.861 | 1.44249 |
| -1.314 | 1.34756 |
| -1.272 | 1.33339 |
| -1.133 | 1.206 |
| -0.924 | 1.09356 |
| -1.002 | 0.673685 |
| -0.932 | 0.663861 |
| -1.001 | 0.650315 |
| -0.86 | 0.520842 |
| -0.816 | 0.429002 |
| -0.739 | 0.331698 |
| -0.728 | 0.303941 |
| -0.705 | 0.26484 |
| -0.705 | 0.261981 |
| -0.7 | 0.260744 |
| -0.709 | 0.20577 |
| -0.57 | 0.166789 |
| -0.455 | 0.0631597 |
| -0.482 | 0.0288309 |
| -0.609 | 0.0280195 |
| -0.374 | 0.0279083 |
| -0.328 | 0.027339 |
| -0.465 | 0.00989212 |
| -0.132 | 0.0013789 |
| -0.338 | 0.000812891 |
| -0.146 | 0.000434512 |
| -0.288 | 0.000391041 |
| -2.03 | 5 |
| 0.551 | 5 |
| 0.644 | 5 |
| -2.028 | 5 |
| 1.699 | 5 |
| -1.936 | 5 |
| -1.202 | 5 |
| -1.389 | 5 |
| 0.418 | 5 |
| 0.373 | 5 |
| 0.234 | 5 |
| 0.687 | 5 |
| -1.411 | 5 |
| 0.669 | 5 |
| 0.718 | 5 |
| 0.27 | 5 |
| 1.303 | 5 |
| 1.465 | 5 |
| 0.19 | 5 |
| 0.738 | 5 |
| 0.349 | 5 |
| 0.14 | 5 |
| -1.861 | 5 |
| 0.43 | 5 |
| 0.214 | 5 |
| -1.085 | 5 |
| 0.778 | 5 |
| -1.733 | 5 |
| 0.063 | 5 |
| 0.264 | 5 |
| 0.108 | 5 |
| 0.277 | 5 |
| 0.595 | 5 |
| 0.003 | 5 |
| 0.196 | 5 |
| 0.309 | 5 |
| -1.173 | 5 |
| -1.148 | 5 |
| 0.492 | 5 |
| 0.251 | 5 |
| 0.052 | 5 |
| 0.282 | 5 |
| 2.244 | 5 |
| 0.634 | 5 |
| -1.398 | 5 |
| -1.794 | 5 |
| -1.705 | 5 |
| 0.383 | 5 |
| 0.416 | 5 |
| 1.02 | 5 |
| 0.256 | 5 |
| -1.685 | 5 |
| 1.616 | 5 |
| 0.501 | 5 |
| -1.55 | 5 |
| 0.537 | 5 |
| 0.06 | 5 |
| 0.19 | 5 |
| 0.16 | 5 |
| 0.273 | 5 |
| 0.343 | 5 |
| 0.612 | 5 |
| -1.532 | 5 |
| 0.543 | 5 |
| 0.689 | 5 |
| 0.341 | 5 |
| -1.346 | 5 |
| 0.987 | 5 |
| -2.12 | 5 |
| -1.988 | 5 |
| 1.712 | 5 |
| 0.209 | 5 |
| 0.727 | 5 |
| 0.147 | 5 |
| -2.1 | 5 |
| -2.395 | 5 |
| 0.081 | 5 |
| 0.22 | 5 |
| 0.247 | 5 |
| 1.133 | 5 |
| 0.757 | 5 |
| -1.105 | 5 |
| 1.21 | 5 |
| -1.972 | 5 |
| -1.927 | 5 |
| 0.25 | 5 |
| 0.919 | 5 |
| 0.371 | 5 |
| 0.244 | 5 |
| -1.623 | 5 |
| -1.875 | 5 |
| 1.084 | 5 |
| -1.741 | 5 |
| -1.919 | 5 |
| -1.374 | 5 |
| -1.262 | 5 |
| -1.79 | 5 |
| -1.225 | 5 |
| -1.127 | 5 |
| 0.369 | 5 |
| 0.356 | 4.69897 |
| 0.203 | 4.69897 |
| 0.467 | 4.69897 |
| 0.274 | 4.69897 |
| 0.201 | 4.69897 |
| 0.414 | 4.69897 |
| 0.061 | 4.69897 |
| -1.475 | 4.69897 |
| -1.622 | 4.69897 |
| -1.193 | 4.69897 |
| 0.168 | 4.69897 |
| 0.331 | 4.69897 |
| 0.782 | 4.69897 |
| 0.594 | 4.69897 |
| 0.42 | 4.69897 |
| -1.443 | 4.69897 |
| -1.635 | 4.69897 |
| 0.21 | 4.69897 |
| 0.231 | 4.69897 |
| 0.469 | 4.69897 |
| 0.414 | 4.69897 |
| 1.216 | 4.69897 |
| -1.387 | 4.69897 |
| -1.741 | 4.69897 |
| 0.321 | 4.69897 |
| -1.643 | 4.69897 |
| -0.853 | 4.69897 |
| 0.408 | 4.69897 |
| 0.122 | 4.69897 |
| 0.261 | 4.69897 |
| 0.368 | 4.52288 |
| 0.114 | 4.52288 |
| 0.091 | 4.52288 |
| 0.251 | 4.52288 |
| 0.663 | 4.52288 |
| 0.188 | 4.52288 |
| -1.698 | 4.52288 |
| 0.13 | 4.52288 |
| 0.192 | 4.39794 |
| 0.556 | 4.39794 |
| -1.931 | 4.39794 |
| -2.2 | 4.39794 |
| -1.061 | 4.39794 |
| 0.424 | 4.39794 |
| 0.315 | 4.39794 |
| 0.087 | 4.39794 |
| -1.589 | 4.39794 |
| 0.25 | 4.39794 |
| -1.494 | 4.30103 |
| 0.441 | 4.30103 |
| 0.117 | 4.30103 |
| 0.02 | 4.22185 |
| 0.154 | 4.22185 |
| -1.742 | 4.22185 |
| 0.292 | 4.22185 |
| 0.162 | 4.22185 |
| -1.313 | 4.22185 |
| -1.06 | 4.22185 |
| 0.457 | 4.1549 |
| 0.075 | 4.1549 |
| 0.369 | 4.1549 |
| 0.023 | 4.1549 |
| 0.255 | 4.09691 |
| -1.171 | 4.09691 |

| Tajima's D | Outlier Sign Taj D |
| --- | --- |
| 0.28 | 3.34679 |
| 0.15 | 3.3098 |
| -1.075 | 3.08619 |

| Tajima's D | Outlier NS Taj D |
| --- | --- |
| 0.086 | 2.86967 |
| -1.68 | 2.83268 |
| 0.151 | 2.79048 |
| -1.356 | 2.44733 |
| -1.565 | 1.89894 |
| -1.031 | 1.55878 |
| -0.861 | 1.44249 |
| -1.314 | 1.34756 |
| -1.272 | 1.33339 |
| -1.133 | 1.206 |
| -0.924 | 1.09356 |
| -1.002 | 0.673685 |
| -0.932 | 0.663861 |
| -1.001 | 0.650315 |
| -0.86 | 0.520842 |
| -0.816 | 0.429002 |
| -0.739 | 0.331698 |
| -0.728 | 0.303941 |
| -0.705 | 0.26484 |
| -0.705 | 0.261981 |
| -0.7 | 0.260744 |
| -0.709 | 0.20577 |
| -0.57 | 0.166789 |
| -0.455 | 0.0631597 |
| -0.482 | 0.0288309 |
| -0.609 | 0.0280195 |
| -0.374 | 0.0279083 |
| -0.328 | 0.027339 |
| -0.465 | 0.00989212 |
| -0.132 | 0.0013789 |
| -0.338 | 0.000812891 |
| -0.146 | 0.000434512 |
| -0.288 | 0.000391041 |

**Figure S3B**

Chiricahua

-Log10(P)

| Tajima's D | All Data |
| --- | --- |
| 0.399 | 5 |
| 1.158 | 4.69897 |
| 1.059 | 4.69897 |
| 1.093 | 4.52288 |
| 0.426 | 4.22185 |
| 0.784 | 3.88606 |
| 1.005 | 3.61979 |
| 0.617 | 2.96257 |
| 0.395 | 2.9393 |
| 0.309 | 2.73518 |
| 0.503 | 2.50724 |
| 0.962 | 2.50585 |
| 0.441 | 2.48017 |
| 0.518 | 2.39903 |
| 0.527 | 2.25337 |
| 0.363 | 2.15304 |
| -1.084 | 2.15304 |
| 0.355 | 1.94654 |
| -0.972 | 1.77754 |
| -0.651 | 1.75424 |
| 0.325 | 1.68719 |
| 0.354 | 1.58186 |
| 0.134 | 1.5812 |
| 0.011 | 1.47939 |
| 0.719 | 1.47006 |
| 0.032 | 1.46686 |
| 0.31 | 1.35932 |
| 0.508 | 1.14893 |
| -0.632 | 0.85608 |
| 0.041 | 0.726004 |
| -0.549 | 0.698775 |
| -0.461 | 0.49447 |
| -0.378 | 0.416665 |
| -0.302 | 0.378439 |
| -0.295 | 0.322749 |
| -0.249 | 0.272743 |
| -0.167 | 0.193494 |
| -0.086 | 0.193284 |
| -0.157 | 0.179766 |
| -0.207 | 0.126075 |
| -1.636 | 5 |
| -1.266 | 5 |
| -1.33 | 5 |
| 0.59 | 5 |
| 1.294 | 5 |
| 0.099 | 5 |
| 1.055 | 5 |
| 1.847 | 5 |
| -1.215 | 5 |
| 0.198 | 5 |
| 2.415 | 5 |
| -1.012 | 5 |
| -0.96 | 5 |
| 0.974 | 5 |
| 0.891 | 5 |
| -1.299 | 5 |
| 2.121 | 5 |
| -1.039 | 5 |
| 0.836 | 5 |
| 0.88 | 5 |
| -0.614 | 5 |
| 0.951 | 5 |
| 1.149 | 5 |
| 1.023 | 5 |
| -0.831 | 5 |
| 1.128 | 5 |
| -0.708 | 5 |
| -1.212 | 5 |
| 0.915 | 5 |
| 0.658 | 5 |
| 0.43 | 5 |
| 1.136 | 5 |
| -1.366 | 5 |
| -1.575 | 5 |
| -1.476 | 5 |
| -1.519 | 5 |
| 0.981 | 5 |
| 1.364 | 5 |
| 1.072 | 5 |
| 1.027 | 5 |
| 0.566 | 5 |
| 0.291 | 5 |
| 0.571 | 5 |
| -1.329 | 5 |
| 0.948 | 5 |
| 0.682 | 5 |
| 1.055 | 5 |
| -1.195 | 5 |
| 0.562 | 5 |
| 0.731 | 5 |
| 0.588 | 5 |
| 0.754 | 5 |
| 1.042 | 5 |
| 1.506 | 5 |
| -0.798 | 5 |
| 1.02 | 5 |
| -0.77 | 5 |
| -1.262 | 5 |
| -0.939 | 5 |
| 0.297 | 5 |
| 0.296 | 5 |
| 0.38 | 5 |
| -1.507 | 5 |
| -1.046 | 5 |
| 1.297 | 5 |
| 0.075 | 5 |
| 0.95 | 5 |
| 0.253 | 5 |
| 1.149 | 5 |
| -0.666 | 5 |
| -0.876 | 5 |
| 0.487 | 5 |
| -1.043 | 5 |
| -1.379 | 5 |
| -1.246 | 5 |
| 0.466 | 5 |
| 0.926 | 5 |
| -0.63 | 5 |
| 0.696 | 5 |
| 1.145 | 5 |
| -0.946 | 5 |
| 0.483 | 5 |
| -1.564 | 5 |
| -1.213 | 5 |
| 1.078 | 5 |
| -1.835 | 5 |
| 0.881 | 5 |
| 0.792 | 5 |
| 0.776 | 5 |
| 1.582 | 5 |
| 1.249 | 5 |
| 0.476 | 5 |
| 0.835 | 5 |
| -1.386 | 5 |
| -0.915 | 5 |
| -1.355 | 5 |
| 1.039 | 5 |
| -0.75 | 5 |
| 1.053 | 5 |
| 0.525 | 5 |
| 0.911 | 5 |
| 1.117 | 5 |
| 1.378 | 5 |
| -1.395 | 5 |
| -1.126 | 5 |
| 1.186 | 5 |
| 0.429 | 5 |
| 0.519 | 5 |
| -1.059 | 5 |
| 0.616 | 5 |
| -0.98 | 5 |
| -1.086 | 5 |
| 1.623 | 5 |
| 0.902 | 5 |
| -0.566 | 5 |
| 0.884 | 5 |
| 0.724 | 5 |
| 0.468 | 5 |
| -1.597 | 5 |
| 0.813 | 5 |
| 0.864 | 5 |
| 1.112 | 5 |
| 0.62 | 5 |
| 0.842 | 5 |
| -0.868 | 5 |
| 1.147 | 5 |
| -1.118 | 5 |
| -0.899 | 5 |
| -1.454 | 5 |
| 1.091 | 5 |
| 0.621 | 5 |
| 0.618 | 5 |
| 1.257 | 5 |
| -0.729 | 5 |
| -1.08 | 5 |
| 0.666 | 5 |
| 0.601 | 5 |
| 1.367 | 5 |
| 0.205 | 5 |
| -1.188 | 5 |
| 0.909 | 5 |
| 0.597 | 5 |
| 0.765 | 5 |
| 0.256 | 5 |
| 1.019 | 5 |
| -1.305 | 5 |
| 1.369 | 5 |
| -1.154 | 5 |
| 1.105 | 5 |
| -1.301 | 5 |
| -0.714 | 5 |
| 1.355 | 5 |
| 0.408 | 5 |
| 0.959 | 5 |
| 1.266 | 5 |
| -0.829 | 5 |
| -1.536 | 5 |
| -1.343 | 5 |
| 0.671 | 5 |
| 0.872 | 5 |

| Tajima's D | Outlier Sign Taj D |
| --- | --- |
| 0.399 | 5 |
| 1.158 | 4.69897 |
| 1.059 | 4.69897 |
| 1.093 | 4.52288 |
| 0.426 | 4.22185 |
| 0.784 | 3.88606 |
| 1.005 | 3.61979 |
| 0.617 | 2.96257 |
| 0.395 | 2.9393 |
| 0.309 | 2.73518 |
| 0.503 | 2.50724 |
| 0.962 | 2.50585 |
| 0.441 | 2.48017 |
| 0.518 | 2.39903 |
| 0.527 | 2.25337 |
| 0.363 | 2.15304 |
| -1.084 | 2.15304 |
| 0.355 | 1.94654 |
| -0.972 | 1.77754 |
| -0.651 | 1.75424 |
| 0.325 | 1.68719 |

| Tajima's D | Outlier NS Taj D |
| --- | --- |
| 0.354 | 1.58186 |
| 0.134 | 1.5812 |
| 0.011 | 1.47939 |
| 0.719 | 1.47006 |
| 0.032 | 1.46686 |
| 0.31 | 1.35932 |
| 0.508 | 1.14893 |
| -0.632 | 0.85608 |
| 0.041 | 0.726004 |
| -0.549 | 0.698775 |
| -0.461 | 0.49447 |
| -0.378 | 0.416665 |
| -0.302 | 0.378439 |
| -0.295 | 0.322749 |
| -0.249 | 0.272743 |
| -0.167 | 0.193494 |
| -0.086 | 0.193284 |
| -0.157 | 0.179766 |
| -0.207 | 0.126075 |

**Figure S3C**

Pikes Peak

-Log10(P)

| Tajima's D | All Data |
| --- | --- |
| -1.928 | 5 |
| 0.902 | 5 |
| -1.48 | 5 |
| -2.034 | 5 |
| 1.005 | 4.69897 |
| -1.473 | 4.69897 |
| 1.531 | 4.52288 |
| -1.289 | 4.22185 |
| -1.401 | 3.74473 |
| -1.127 | 3.72125 |
| -0.921 | 3.72125 |
| -0.633 | 3.55284 |
| 0.974 | 3.16749 |
| -1.176 | 3.11351 |
| 0.289 | 3.04096 |
| -0.568 | 3.02228 |
| 0.037 | 2.96257 |
| -0.767 | 2.90309 |
| -1.053 | 2.88941 |
| -0.814 | 2.73518 |
| -0.468 | 2.66354 |
| -0.963 | 2.64975 |
| 0.574 | 2.57512 |
| -1.229 | 2.54516 |
| -1.326 | 2.53018 |
| -1.203 | 2.51999 |
| 0.623 | 2.41454 |
| 0.529 | 2.26043 |
| -1.011 | 2.12843 |
| 0.187 | 2.07366 |
| -0.763 | 2.03527 |
| -0.626 | 2.02733 |
| -1.151 | 2.02457 |
| 0.63 | 1.98758 |
| 0.08 | 1.94424 |
| -0.606 | 1.8548 |
| -0.867 | 1.82362 |
| -1.098 | 1.81361 |
| -0.795 | 1.80438 |
| -0.698 | 1.80244 |
| -0.999 | 1.73898 |
| -1.027 | 1.67572 |
| -0.859 | 1.65228 |
| -0.491 | 1.63358 |
| 0.518 | 1.61422 |
| 0.342 | 1.59585 |
| -1.073 | 1.59142 |
| 0.019 | 1.59125 |
| -0.341 | 1.57561 |
| 0.228 | 1.50473 |
| 0.172 | 1.47782 |
| -0.795 | 1.47224 |
| -0.346 | 1.44382 |
| -0.689 | 1.3026 |
| -0.509 | 1.28508 |
| -0.375 | 1.27572 |
| 0.188 | 1.25688 |
| 0.121 | 1.25508 |
| -0.327 | 1.23567 |
| 0.695 | 1.23025 |
| 0.143 | 1.22702 |
| 0.518 | 1.20986 |
| 0.64 | 1.2067 |
| 0.154 | 1.20094 |
| -0.769 | 1.19511 |
| -0.611 | 1.18436 |
| -0.53 | 1.14972 |
| 0.021 | 1.14557 |
| 0.351 | 1.14261 |
| 0.143 | 1.09469 |
| -0.434 | 1.08276 |
| 0.479 | 1.07753 |
| -0.763 | 1.06656 |
| 0.221 | 1.06053 |
| -0.69 | 1.04354 |
| -0.741 | 1.03626 |
| -0.341 | 1.01439 |
| -0.764 | 1.01399 |
| -0.468 | 1.01291 |
| -0.322 | 0.980468 |
| 0.546 | 0.962693 |
| 0.239 | 0.935767 |
| -0.368 | 0.926795 |
| 0.429 | 0.919843 |
| -0.766 | 0.905494 |
| -0.39 | 0.888637 |
| -0.337 | 0.886591 |
| -0.433 | 0.862773 |
| -0.624 | 0.808018 |
| -0.643 | 0.80482 |
| 0.091 | 0.802802 |
| 0.146 | 0.772113 |
| -0.551 | 0.721841 |
| -0.355 | 0.714645 |
| 0.22 | 0.708254 |
| -0.288 | 0.699361 |
| 0.167 | 0.699187 |
| -0.427 | 0.69715 |
| -0.385 | 0.686935 |
| 0.116 | 0.638726 |
| 0.171 | 0.634381 |
| -0.367 | 0.613376 |
| -0.408 | 0.595354 |
| 0.053 | 0.580573 |
| 0.018 | 0.575102 |
| -0.384 | 0.566742 |
| -0.394 | 0.545689 |
| 0.027 | 0.47843 |
| 0.111 | 0.465987 |
| -0.345 | 0.43348 |
| -0.355 | 0.431528 |
| -0.445 | 0.431024 |
| -0.347 | 0.413919 |
| -0.242 | 0.357763 |
| -0.22 | 0.35637 |
| -0.244 | 0.351786 |
| -0.24 | 0.347058 |
| -0.28 | 0.345342 |
| -0.213 | 0.33208 |
| -0.211 | 0.308697 |
| -0.196 | 0.296056 |
| -0.201 | 0.287569 |
| -0.188 | 0.279163 |
| -0.196 | 0.270074 |
| -0.174 | 0.25877 |
| -0.176 | 0.257573 |
| -0.165 | 0.241398 |
| -0.139 | 0.231725 |
| -0.144 | 0.227634 |
| -0.066 | 0.226909 |
| -0.148 | 0.222015 |
| -0.155 | 0.220822 |
| -0.082 | 0.216826 |
| -0.137 | 0.210321 |
| -0.145 | 0.205944 |
| -0.032 | 0.179937 |
| -0.071 | 0.168175 |
| -0.01 | 0.157216 |
| -0.16 | 0.154697 |
| -0.08 | 0.153508 |
| -0.104 | 0.146466 |
| -0.272 | 0.143017 |
| -0.143 | 0.125959 |
| -0.072 | 0.116316 |
| -0.086 | 0.115669 |
| -0.008 | 0.106999 |
| -0.04 | 0.0950303 |
| -0.074 | 0.0906366 |
| -0.001 | 0.0145466 |
| 0.17 | 5 |
| 1.694 | 5 |
| -1.273 | 5 |
| -1.603 | 5 |
| 1.392 | 5 |
| -1.32 | 5 |
| 1.108 | 5 |
| -1.67 | 5 |
| 1.343 | 5 |
| 0.64 | 5 |
| 1.047 | 5 |
| -1.551 | 5 |
| -0.65 | 5 |
| -1.309 | 5 |
| 0.388 | 5 |
| 0.681 | 5 |
| 1.381 | 5 |
| -1.183 | 5 |
| 0.191 | 5 |
| 1.324 | 5 |
| 1.513 | 5 |
| 1.191 | 5 |
| -1.63 | 5 |
| 1.027 | 5 |
| 0.836 | 5 |
| 0.851 | 5 |
| 0.71 | 5 |
| 1.196 | 5 |
| 0.971 | 5 |
| 0.637 | 5 |
| -0.809 | 5 |
| 1.168 | 5 |
| -1.452 | 5 |
| 1.203 | 5 |
| 1.024 | 5 |
| 1.144 | 5 |
| 0.926 | 5 |
| 1.112 | 5 |
| -0.892 | 5 |
| 0.724 | 5 |
| 0.992 | 5 |
| -1.263 | 5 |
| -1.802 | 5 |
| -1.728 | 5 |
| -1.618 | 5 |
| 1.492 | 5 |
| -1.359 | 5 |
| 1.219 | 5 |
| -1.673 | 5 |
| -1.101 | 5 |
| 0.815 | 5 |

| Tajima's D | Outlier Sign Tajima D |
| --- | --- |
| -1.928 | 5 |
| 0.902 | 5 |
| -1.48 | 5 |
| -2.034 | 5 |
| 1.005 | 4.69897 |
| -1.473 | 4.69897 |
| 1.531 | 4.52288 |
| -1.289 | 4.22185 |
| -1.401 | 3.74473 |
| -1.127 | 3.72125 |
| -0.921 | 3.72125 |
| -0.633 | 3.55284 |
| 0.974 | 3.16749 |
| -1.176 | 3.11351 |
| 0.289 | 3.04096 |
| -0.568 | 3.02228 |
| 0.037 | 2.96257 |
| -0.767 | 2.90309 |
| -1.053 | 2.88941 |
| -0.814 | 2.73518 |
| -0.468 | 2.66354 |
| -0.963 | 2.64975 |
| 0.574 | 2.57512 |
| -1.229 | 2.54516 |
| -1.326 | 2.53018 |
| -1.203 | 2.51999 |
| 0.623 | 2.41454 |
| 0.529 | 2.26043 |
| -1.011 | 2.12843 |
| 0.187 | 2.07366 |
| -0.763 | 2.03527 |
| -0.626 | 2.02733 |
| -1.151 | 2.02457 |
| 0.63 | 1.98758 |
| 0.08 | 1.94424 |
| -0.606 | 1.8548 |
| -0.867 | 1.82362 |
| -1.098 | 1.81361 |
| -0.795 | 1.80438 |
| -0.698 | 1.80244 |
| -0.999 | 1.73898 |
| -1.027 | 1.67572 |

| Tajima's D | Outlier NS Tajima D |
| --- | --- |
| -0.859 | 1.65228 |
| -0.491 | 1.63358 |
| 0.518 | 1.61422 |
| 0.342 | 1.59585 |
| -1.073 | 1.59142 |
| 0.019 | 1.59125 |
| -0.341 | 1.57561 |
| 0.228 | 1.50473 |
| 0.172 | 1.47782 |
| -0.795 | 1.47224 |
| -0.346 | 1.44382 |
| -0.689 | 1.3026 |
| -0.509 | 1.28508 |
| -0.375 | 1.27572 |
| 0.188 | 1.25688 |
| 0.121 | 1.25508 |
| -0.327 | 1.23567 |
| 0.695 | 1.23025 |
| 0.143 | 1.22702 |
| 0.518 | 1.20986 |
| 0.64 | 1.2067 |
| 0.154 | 1.20094 |
| -0.769 | 1.19511 |
| -0.611 | 1.18436 |
| -0.53 | 1.14972 |
| 0.021 | 1.14557 |
| 0.351 | 1.14261 |
| 0.143 | 1.09469 |
| -0.434 | 1.08276 |
| 0.479 | 1.07753 |
| -0.763 | 1.06656 |
| 0.221 | 1.06053 |
| -0.69 | 1.04354 |
| -0.741 | 1.03626 |
| -0.341 | 1.01439 |
| -0.764 | 1.01399 |
| -0.468 | 1.01291 |
| -0.322 | 0.980468 |
| 0.546 | 0.962693 |
| 0.239 | 0.935767 |
| -0.368 | 0.926795 |
| 0.429 | 0.919843 |
| -0.766 | 0.905494 |
| -0.39 | 0.888637 |
| -0.337 | 0.886591 |
| -0.433 | 0.862773 |
| -0.624 | 0.808018 |
| -0.643 | 0.80482 |
| 0.091 | 0.802802 |
| 0.146 | 0.772113 |
| -0.551 | 0.721841 |
| -0.355 | 0.714645 |
| 0.22 | 0.708254 |
| -0.288 | 0.699361 |
| 0.167 | 0.699187 |
| -0.427 | 0.69715 |
| -0.385 | 0.686935 |
| 0.116 | 0.638726 |
| 0.171 | 0.634381 |
| -0.367 | 0.613376 |
| -0.408 | 0.595354 |
| 0.053 | 0.580573 |
| 0.018 | 0.575102 |
| -0.384 | 0.566742 |
| -0.394 | 0.545689 |
| 0.027 | 0.47843 |
| 0.111 | 0.465987 |
| -0.345 | 0.43348 |
| -0.355 | 0.431528 |
| -0.445 | 0.431024 |
| -0.347 | 0.413919 |
| -0.242 | 0.357763 |
| -0.22 | 0.35637 |
| -0.244 | 0.351786 |
| -0.24 | 0.347058 |
| -0.28 | 0.345342 |
| -0.213 | 0.33208 |
| -0.211 | 0.308697 |
| -0.196 | 0.296056 |
| -0.201 | 0.287569 |
| -0.188 | 0.279163 |
| -0.196 | 0.270074 |
| -0.174 | 0.25877 |
| -0.176 | 0.257573 |
| -0.165 | 0.241398 |
| -0.139 | 0.231725 |
| -0.144 | 0.227634 |
| -0.066 | 0.226909 |
| -0.148 | 0.222015 |
| -0.155 | 0.220822 |
| -0.082 | 0.216826 |
| -0.137 | 0.210321 |
| -0.145 | 0.205944 |
| -0.032 | 0.179937 |
| -0.071 | 0.168175 |
| -0.01 | 0.157216 |
| -0.16 | 0.154697 |
| -0.08 | 0.153508 |
| -0.104 | 0.146466 |
| -0.272 | 0.143017 |
| -0.143 | 0.125959 |
| -0.072 | 0.116316 |
| -0.086 | 0.115669 |
| -0.008 | 0.106999 |
| -0.04 | 0.0950303 |
| -0.074 | 0.0906366 |
| -0.001 | 0.0145466 |

**Figure S3D**

Standard

-Log10(P)

| Tajima's D | All Data |
| --- | --- |
| -0.604 | 5 |
| -1.137 | 5 |
| 1.083 | 5 |
| 1.613 | 5 |
| 1.707 | 5 |
| 1.368 | 5 |
| -1.157 | 5 |
| -1.464 | 4.52288 |
| -1.226 | 4.30103 |
| 0.416 | 4.22185 |
| -1.125 | 3.3098 |
| 0.776 | 2.89279 |
| -0.787 | 2.87943 |
| -1.091 | 2.84466 |
| -1.389 | 2.82974 |
| -0.884 | 2.68403 |
| -1.378 | 2.37263 |
| 0.75 | 2.35458 |
| -0.733 | 2.32331 |
| -1.286 | 2.29671 |
| -1.317 | 2.27819 |
| 0.167 | 2.17005 |
| 0.638 | 2.13253 |
| 0.492 | 1.99055 |
| -0.783 | 1.99055 |
| 1.285 | 1.96337 |
| -0.962 | 1.84894 |
| 0.651 | 1.6332 |
| 0.359 | 1.61654 |
| 0.217 | 1.61618 |
| -0.903 | 1.59636 |
| -0.505 | 1.58737 |
| 0.555 | 1.58453 |
| 0.275 | 1.57365 |
| -0.902 | 1.52244 |
| -0.862 | 1.42493 |
| -0.476 | 1.36694 |
| -0.69 | 1.23448 |
| -0.734 | 1.18936 |
| -0.399 | 1.18936 |
| -0.844 | 1.16368 |
| -0.573 | 1.10869 |
| 0.56 | 1.07335 |
| -0.286 | 1.07166 |
| -0.553 | 1.0348 |
| 0.29 | 0.856361 |
| -0.313 | 0.84952 |
| 0.166 | 0.827485 |
| -0.47 | 0.812649 |
| 0.365 | 0.790914 |
| -0.559 | 0.764143 |
| 0.133 | 0.759376 |
| -1.175 | 0.749848 |
| -0.501 | 0.747171 |
| 0.327 | 0.745839 |
| -0.334 | 0.696502 |
| 0.161 | 0.677305 |
| 0.168 | 0.639482 |
| -0.325 | 0.603591 |
| -0.296 | 0.597257 |
| 0.022 | 0.592116 |
| 0.122 | 0.581699 |
| 0.096 | 0.540095 |
| 0.135 | 0.516513 |
| -0.311 | 0.483941 |
| 0.02 | 0.482066 |
| -0.191 | 0.351074 |
| -0.189 | 0.334719 |
| -0.375 | 0.307426 |
| -0.154 | 0.304186 |
| -0.145 | 0.290247 |
| -0.129 | 0.267896 |
| -0.327 | 0.23046 |
| -0.041 | 0.208365 |
| -0.052 | 0.161535 |
| -0.265 | 0.155007 |
| -0.009 | 0.115182 |
| -0.041 | 0.106133 |
| -0.039 | 0.102505 |
| -0.218 | 0.06584 |
| -0.207 | 0.056738 |
| 0.644 | 5 |
| 1.228 | 5 |
| 0.38 | 5 |
| -1.586 | 5 |
| 0.917 | 5 |
| 0.736 | 5 |
| 0.744 | 5 |
| 0.668 | 5 |
| 0.671 | 5 |
| -0.782 | 5 |
| 1.617 | 5 |
| 0.541 | 5 |
| -1.485 | 5 |
| 1.079 | 5 |
| -1.453 | 5 |
| -1.528 | 5 |
| -1.219 | 5 |
| 2.27 | 5 |
| -0.544 | 5 |
| -1.351 | 5 |
| 1.042 | 5 |
| 1.151 | 5 |
| 0.671 | 5 |
| 1.409 | 5 |
| 1.237 | 5 |
| 1.625 | 5 |
| -0.798 | 5 |
| 1.877 | 5 |
| 1.462 | 5 |
| 0.878 | 5 |
| 0.675 | 5 |
| 1.025 | 5 |
| -1.423 | 5 |
| -1.251 | 5 |
| 0.145 | 5 |
| -1.655 | 5 |
| -1.589 | 5 |
| -1.213 | 5 |
| -1.358 | 5 |
| -1.752 | 5 |
| -1.331 | 5 |
| -0.757 | 5 |
| 0.881 | 5 |
| -0.851 | 5 |
| -1.366 | 5 |
| 1.206 | 5 |
| 1.462 | 5 |
| 0.4 | 5 |
| 1.128 | 5 |
| 0.961 | 5 |
| -1.688 | 5 |
| 0.843 | 5 |
| 1.394 | 5 |
| 1.586 | 5 |
| -1.19 | 5 |
| 1.512 | 5 |
| -0.737 | 5 |
| -1.278 | 5 |
| -1.287 | 5 |
| 0.535 | 5 |
| 1.198 | 5 |
| -1.574 | 5 |
| 0.864 | 5 |
| 1.262 | 5 |
| -1.565 | 5 |
| 0.374 | 5 |
| -1.044 | 5 |
| 0.883 | 5 |
| 0.956 | 5 |
| -0.611 | 5 |
| -1.372 | 5 |
| 0.674 | 5 |
| -1.515 | 5 |
| -1.534 | 5 |
| 0.952 | 5 |
| 0.822 | 5 |
| -1.296 | 5 |
| -1.006 | 5 |
| -1.379 | 5 |
| -1.521 | 5 |
| -1.559 | 5 |
| 1.577 | 5 |
| -1.481 | 5 |
| -1.21 | 5 |
| -1.277 | 5 |
| 0.711 | 5 |
| 0.657 | 5 |
| 0.294 | 5 |
| 0.808 | 5 |
| -1.14 | 5 |
| -1.554 | 5 |
| -1.206 | 5 |
| 0.698 | 5 |
| 0.417 | 5 |
| 0.753 | 5 |
| 0.889 | 5 |
| 0.832 | 5 |
| -1.327 | 5 |
| 0.074 | 5 |
| 0.985 | 5 |
| -0.924 | 5 |
| -1.312 | 5 |
| 0.95 | 5 |
| 1.118 | 5 |
| 2.07 | 5 |
| 0.858 | 5 |
| -1.316 | 5 |
| 0.878 | 5 |
| -0.974 | 5 |
| 1.023 | 5 |
| 1.409 | 5 |
| 0.84 | 5 |
| -0.929 | 5 |
| 0.779 | 5 |
| 0.819 | 5 |
| -0.721 | 5 |
| 1.417 | 5 |
| -1.698 | 5 |

| Tajima's D | Outlier Sign Taj D |
| --- | --- |
| -0.604 | 5 |
| -1.137 | 5 |
| 1.083 | 5 |
| 1.613 | 5 |
| 1.707 | 5 |
| 1.368 | 5 |
| -1.157 | 5 |
| -1.464 | 4.52288 |
| -1.226 | 4.30103 |
| 0.416 | 4.22185 |
| -1.125 | 3.3098 |
| 0.776 | 2.89279 |
| -0.787 | 2.87943 |
| -1.091 | 2.84466 |
| -1.389 | 2.82974 |
| -0.884 | 2.68403 |
| -1.378 | 2.37263 |
| 0.75 | 2.35458 |
| -0.733 | 2.32331 |
| -1.286 | 2.29671 |
| -1.317 | 2.27819 |
| 0.167 | 2.17005 |
| 0.638 | 2.13253 |
| 0.492 | 1.99055 |
| -0.783 | 1.99055 |
| 1.285 | 1.96337 |

| Tajima's D | Outlier NS Taj D |
| --- | --- |
| -0.962 | 1.84894 |
| 0.651 | 1.6332 |
| 0.359 | 1.61654 |
| 0.217 | 1.61618 |
| -0.903 | 1.59636 |
| -0.505 | 1.58737 |
| 0.555 | 1.58453 |
| 0.275 | 1.57365 |
| -0.902 | 1.52244 |
| -0.862 | 1.42493 |
| -0.476 | 1.36694 |
| -0.69 | 1.23448 |
| -0.734 | 1.18936 |
| -0.399 | 1.18936 |
| -0.844 | 1.16368 |
| -0.573 | 1.10869 |
| 0.56 | 1.07335 |
| -0.286 | 1.07166 |
| -0.553 | 1.0348 |
| 0.29 | 0.856361 |
| -0.313 | 0.84952 |
| 0.166 | 0.827485 |
| -0.47 | 0.812649 |
| 0.365 | 0.790914 |
| -0.559 | 0.764143 |
| 0.133 | 0.759376 |
| -1.175 | 0.749848 |
| -0.501 | 0.747171 |
| 0.327 | 0.745839 |
| -0.334 | 0.696502 |
| 0.161 | 0.677305 |
| 0.168 | 0.639482 |
| -0.325 | 0.603591 |
| -0.296 | 0.597257 |
| 0.022 | 0.592116 |
| 0.122 | 0.581699 |
| 0.096 | 0.540095 |
| 0.135 | 0.516513 |
| -0.311 | 0.483941 |
| 0.02 | 0.482066 |
| -0.191 | 0.351074 |
| -0.189 | 0.334719 |
| -0.375 | 0.307426 |
| -0.154 | 0.304186 |
| -0.145 | 0.290247 |
| -0.129 | 0.267896 |
| -0.327 | 0.23046 |
| -0.041 | 0.208365 |
| -0.052 | 0.161535 |
| -0.265 | 0.155007 |
| -0.009 | 0.115182 |
| -0.041 | 0.106133 |
| -0.039 | 0.102505 |
| -0.218 | 0.06584 |
| -0.207 | 0.056738 |

**Figure S3E**

Tree Line

-Log10(P)

| Tajima's D | All Data |
| --- | --- |
| 0.595 | 4.69897 |
| -1.557 | 4.69897 |
| -1.211 | 4.69897 |
| 0.931 | 4.52288 |
| -0.587 | 4.39794 |
| -0.947 | 4.39794 |
| 0.566 | 4 |
| -1.119 | 3.22915 |
| 0.805 | 2.81531 |
| 0.424 | 2.46471 |
| -0.76 | 2.35067 |
| -0.665 | 2.14874 |
| -0.878 | 2.07212 |
| -0.653 | 2.00877 |
| -0.552 | 1.8687 |
| -0.863 | 1.84013 |
| -0.681 | 1.72909 |
| -1.152 | 1.56431 |
| 0.251 | 1.38342 |
| 0.379 | 1.35605 |
| -0.62 | 1.22599 |
| -0.331 | 1.20121 |
| -0.546 | 1.18776 |
| -0.532 | 1.15546 |
| 0.349 | 1.01823 |
| -0.509 | 0.997618 |
| -0.336 | 0.951947 |
| -0.412 | 0.799861 |
| -0.53 | 0.750777 |
| 0.206 | 0.738618 |
| -0.498 | 0.722391 |
| 0.041 | 0.580028 |
| -0.359 | 0.577574 |
| -0.338 | 0.567351 |
| -0.258 | 0.552749 |
| -0.307 | 0.521361 |
| -0.446 | 0.50703 |
| -0.327 | 0.506361 |
| -0.363 | 0.486343 |
| -0.278 | 0.452004 |
| -0.307 | 0.446578 |
| -0.231 | 0.390907 |
| -0.195 | 0.338216 |
| -0.27 | 0.318659 |
| -0.192 | 0.305799 |
| -0.188 | 0.300952 |
| -0.175 | 0.285268 |
| -0.164 | 0.273501 |
| -0.104 | 0.178139 |
| -0.155 | 0.161964 |
| -0.044 | 0.154666 |
| -0.122 | 0.131015 |
| -0.122 | 0.0749872 |
| -0.109 | 0.065739 |
| 0.607 | 5 |
| 0.812 | 5 |
| 0.415 | 5 |
| -1.475 | 5 |
| -1.002 | 5 |
| -0.894 | 5 |
| 0.439 | 5 |
| 0.442 | 5 |
| 0.328 | 5 |
| -1.431 | 5 |
| 0.336 | 5 |
| 0.45 | 5 |
| -0.629 | 5 |
| 0.685 | 5 |
| 0.893 | 5 |
| 0.305 | 5 |
| 0.737 | 5 |
| 0.646 | 5 |
| 0.63 | 5 |
| 0.28 | 5 |
| 1.625 | 5 |
| 1.198 | 5 |
| 0.177 | 5 |
| -1.779 | 5 |
| 0.814 | 5 |
| 0.491 | 5 |
| -1.353 | 5 |
| 0.204 | 5 |
| 0.431 | 5 |
| -0.761 | 5 |
| -1.867 | 5 |
| 1.168 | 5 |
| 0.948 | 5 |
| 0.175 | 5 |
| -1.327 | 5 |
| -1.661 | 5 |
| 0.489 | 5 |
| -1 | 5 |
| 0.454 | 5 |
| 0.608 | 5 |
| -1.633 | 5 |
| 0.358 | 5 |
| -1.303 | 5 |
| 0.228 | 5 |
| -0.776 | 5 |
| -0.912 | 5 |
| 0.631 | 5 |
| 0.487 | 5 |
| 0.055 | 5 |
| 0.699 | 5 |
| -1.217 | 5 |
| 0.665 | 5 |
| -0.782 | 5 |
| 0.166 | 5 |
| 0.344 | 5 |
| 0.391 | 5 |
| -1.257 | 5 |
| 1.725 | 5 |
| 1.037 | 5 |
| -1.242 | 5 |
| 1.213 | 5 |
| 1.878 | 5 |
| 0.345 | 5 |
| 0.719 | 5 |
| 0.586 | 5 |
| 0.83 | 5 |
| -1.548 | 5 |
| 0.635 | 5 |
| 0.422 | 5 |
| -0.961 | 5 |
| -0.71 | 5 |
| 0.175 | 5 |
| -0.834 | 5 |
| 0.59 | 5 |
| -0.705 | 5 |
| -0.308 | 5 |
| -1.738 | 5 |
| 0.076 | 5 |
| 1.328 | 5 |
| 0.403 | 5 |
| 0.587 | 5 |
| 0.407 | 5 |
| 0.635 | 5 |
| 1.144 | 5 |
| 0.638 | 5 |
| -1.72 | 5 |
| -1.206 | 5 |
| -0.95 | 5 |
| -0.891 | 5 |
| 0.92 | 5 |
| -1.147 | 5 |
| -1.44 | 5 |
| -1.146 | 5 |
| -1.322 | 5 |
| 0.076 | 5 |
| 1.005 | 5 |
| 1.671 | 5 |
| 0.345 | 5 |
| -1.109 | 5 |
| -1.552 | 5 |
| 1.066 | 5 |
| 0.93 | 5 |
| -1.28 | 5 |
| 0.721 | 5 |
| 1.365 | 5 |
| 0.889 | 5 |
| -0.447 | 5 |
| 1.057 | 5 |
| 0.848 | 5 |
| 0.556 | 5 |
| -1.223 | 5 |
| 0.153 | 5 |
| -1.21 | 5 |
| 0.23 | 5 |
| -1.251 | 4.69897 |
| 1.152 | 4.69897 |
| 0.567 | 4.69897 |
| 0.658 | 4.69897 |
| -1.075 | 4.69897 |
| 0.377 | 4.69897 |
| 0.211 | 4.69897 |
| 0.302 | 4.69897 |
| -1.087 | 4.69897 |
| -1.123 | 4.69897 |
| 0.82 | 4.69897 |
| 0.441 | 4.69897 |
| 0.998 | 4.69897 |
| 1.16 | 4.69897 |
| 0.75 | 4.69897 |
| 0.223 | 4.52288 |
| 0.696 | 4.52288 |
| -0.956 | 4.52288 |
| 0.699 | 4.52288 |
| -1.093 | 4.52288 |
| 0.455 | 4.52288 |
| 0.562 | 4.52288 |
| 0.72 | 4.52288 |
| 0.311 | 4.52288 |
| 1.13 | 4.52288 |
| 0.174 | 4.52288 |
| 0.062 | 4.52288 |
| 1.384 | 4.52288 |
| 0.811 | 4.52288 |
| -1.003 | 4.52288 |
| 0.669 | 4.39794 |
| -1.156 | 4.39794 |

| Tajima's D | Outlier Sign Taj D |
| --- | --- |
| 0.595 | 4.69897 |
| -1.557 | 4.69897 |
| -1.211 | 4.69897 |
| 0.931 | 4.52288 |
| -0.587 | 4.39794 |
| -0.947 | 4.39794 |
| 0.566 | 4 |
| -1.119 | 3.22915 |
| 0.805 | 2.81531 |

| Tajima's D | Outlier NS Taj D |
| --- | --- |
| 0.424 | 2.46471 |
| -0.76 | 2.35067 |
| -0.665 | 2.14874 |
| -0.878 | 2.07212 |
| -0.653 | 2.00877 |
| -0.552 | 1.8687 |
| -0.863 | 1.84013 |
| -0.681 | 1.72909 |
| -1.152 | 1.56431 |
| 0.251 | 1.38342 |
| 0.379 | 1.35605 |
| -0.62 | 1.22599 |
| -0.331 | 1.20121 |
| -0.546 | 1.18776 |
| -0.532 | 1.15546 |
| 0.349 | 1.01823 |
| -0.509 | 0.997618 |
| -0.336 | 0.951947 |
| -0.412 | 0.799861 |
| -0.53 | 0.750777 |
| 0.206 | 0.738618 |
| -0.498 | 0.722391 |
| 0.041 | 0.580028 |
| -0.359 | 0.577574 |
| -0.338 | 0.567351 |
| -0.258 | 0.552749 |
| -0.307 | 0.521361 |
| -0.446 | 0.50703 |
| -0.327 | 0.506361 |
| -0.363 | 0.486343 |
| -0.278 | 0.452004 |
| -0.307 | 0.446578 |
| -0.231 | 0.390907 |
| -0.195 | 0.338216 |
| -0.27 | 0.318659 |
| -0.192 | 0.305799 |
| -0.188 | 0.300952 |
| -0.175 | 0.285268 |
| -0.164 | 0.273501 |
| -0.104 | 0.178139 |
| -0.155 | 0.161964 |
| -0.044 | 0.154666 |
| -0.122 | 0.131015 |
| -0.122 | 0.0749872 |
| -0.109 | 0.065739 |

Figure S3. Plot of Tajima’s D versus -Log2(P) for between 2663 to 2666 genes on Muller C of *D. pseudoobscura*. Outliers with a significantly negative or positive Tajima’s D values are indicated with a pink dot while outliers with non-significant Tajima’s D values are indicated with blue dots. Tajima’s D values for non-outlier genes are indicated with gray dots. A. Arrowhead; B. Chiricahua; C. Pikes Peak; D. Standard; E. Tree Line.

**Table S2.** Chi-Square test of homogeneity of non-outlier and outlier genes in different gene conversion classes for the Arrowhead gene arrangement. Gene conversion (GC) coverage classes are defined by minimum and maximum values from random permutation of gene conversion tracts.

| **GC Coverage Class** | **Non-Outlier** | **Outlier** |
| --- | --- | --- |
| **Below the Minimum**  Coverage < 0.679  Obs  Exp | 944  964.8 | 34  13.2 |
| **Between the Minimum and Maximum**  0.679 <Coverage<2.652  Obs  Exp | 726  718.2 | 2  9.8 |
| **Above Maximum**  Coverage >2.652  Obs  Exp | 962  949.0 | 0  13.0 |
| X^2^ df=2 | 52.7 |  |
| Probability | 3.57 x 10^-12^ |  |

**Table S3.** Chi-Square test of homogeneity of non-outlier and outlier genes in different gene conversion classes for the Chiricahua gene arrangement. Gene conversion (GC) coverage classes are defined by minimum and maximum values from random permutation of gene conversion tracts.

| **Coverage Class** | **Non-Outlier** | **Outlier** |
| --- | --- | --- |
| **Below the Minimum**  Coverage < 0.483  Obs  Exp | 893  914.1 | 35  13.9 |
| **Between the Minimum and Maximum**  0.483 <Coverage<1.735  Obs  Exp | 652  647.1 | 5  9.9 |
| **Above the Maximum**  Coverage >1.735  Obs  Exp | 1083  1066.8 | 0  16.2 |
| X^2^ df=2 | 51.4 |  |
| Probability | 7.05 x 10^-12^ |  |

**Table S4.** Chi-Square test of homogeneity of non-outlier and outlier genes in different gene conversion classes for the Pikes Peak gene arrangement. Gene conversion (GC) coverage classes are defined by minimum and maximum values from random permutation of gene conversion tracts.

| **GC Coverage Class** | **Non-Outlier** | **Outlier** |
| --- | --- | --- |
| **Below the Minimum**  Coverage < 0.233  Obs  Exp | 1254  1314.3 | 138  77.7 |
| **Between the Minimum and Maximum**  0.233 <Coverage<0.922  Obs  Exp | 376  364.4 | 10  21.6 |
| **Above the Maximum**  Coverage >0.922  Obs  Exp | 889  840.3 | 1  49.7 |
| X^2^ df=2 | 106.6 |  |
| Probability | 7.17 x 10^-24^ |  |

**Table S5.** Chi-Square test of homogeneity of non-outlier and outlier genes in different gene conversion classes for the Standard gene arrangement. Gene conversion (GC) coverage classes are defined by minimum and maximum values from random permutation of gene conversion tracts.

| **Coverage Class** | **Non-Outlier** | **Outlier** |
| --- | --- | --- |
| **Below the Minimum**  Coverage < 0.563  Obs  Exp | 1028  1072.0 | 78  34.0 |
| **Between the Minimum and Maximum**  0.563 <Coverage<1.758  Obs  Exp | 646  630.0 | 4  20.0 |
| **Above the Maximum**  Coverage >1.758  Obs  Exp | 912  884.0 | 0  28.0 |
| X^2^ df=2 | 100.9 |  |
| Probability | 1.24 x 10^-22^ |  |

**Table S6**. Chi-Square test of homogeneity of non-outlier and outlier genes in different gene conversion classes for the Tree gene arrangement. Gene conversion (GC) coverage classes are defined by minimum and maximum values from random permutation of gene conversion tracts.

| **Coverage Class** | **Non-Outlier** | **Outlier** |
| --- | --- | --- |
| **Below the Minimum**  Coverage < 0.216  Obs  Exp | 908  936.7 | 48  19.3 |
| **Between the Minimum and the Maximum**  0.216 <Coverage<1.290  Obs  Exp | 741  730.9.0 | 5  15.1 |
| **Above the Maximum**  Coverage >1.290  Obs  Exp | 965  946.4.0 | 1  19.6 |
| X^2^ df=2 | 68.2 |  |
| Probability | 1.58 x 10^-15^ |  |
